# Nephrotoxicity of Immune Checkpoint Inhibitors in Mice with a Human Immune System

**DOI:** 10.64898/2026.05.07.723340

**Authors:** Sarah Asby, Xia Wen, Michael Goedken, Brianna Ames, Shams Shams, Lauren E. Thompson, Jordi Lanis, Zander Kostka-Newman, Kristina Larsen, Scott Tilden, Julie Lang, Lauren M. Aleksunes, Melanie S. Joy

## Abstract

**Introduction:** Immune checkpoint inhibitors (ICIs) enhance antitumor responses by blocking inhibitory receptors, including PD-1 and CTLA-4. Overactivation can trigger systemic toxicity akin to autoimmune diseases, including kidney manifestations. We sought to 1) profile immune signaling and 2) interrogate potential mechanisms of ICI-related kidney injury in a Human Immune System (HIS) tumor-bearing mouse model treated with nivolumab and ipilimumab.

**Methods:** Immunodeficient BRGS (BALB/c-*Rag2^null^Il2rγ^null^Sirpα*^NOD^) neonates were engrafted with human CD34+ cells to generate HIS-BRGS mice. Human MDA-MB-231 tumor cells were implanted subcutaneously; once tumors reached ∼150 mm³, mice received weekly intraperitoneal vehicle (PBS) or ICI (nivolumab 20 mg/kg + ipilimumab 10 mg/kg) for 4 weeks (Veh BRGS n=4; ICI BRGS n=6; Veh HIS-BRGS n=7; ICI HIS-BRGS n=7). Kidneys were evaluated by histopathology (H&E, TEM), flow cytometry for human immune phenotypes, multiplex ELISA (80 human proteins; 10 injury biomarkers), bulk RNA sequencing, and targeted qPCR. Pearson correlations identified predictors of histopathological injury.

**Results:** Renal vasculitis and interstitial nephritis were observed only in ICI-treated HIS-BRGS mice. These kidneys showed a shift toward CD4+ T-cell enrichment with an increased TNF-α production capacity compared to CD8+ counterparts. Toxicity was accompanied by increased renal concentrations of human cytokines, chemokines, and soluble receptors. ICI treatment significantly elevated serine proteases (Granzyme A/B) and NGF-β, while decreasing IL-4. Interstitial nephritis correlated with renal PD-1 and MIF. Renal vasculitis correlated with kidney PD-1, CCL1, MIF, Granzyme A, IL-15, and BAFF. Traditional injury biomarkers (KIM-1, NGAL) remained unchanged; however, a trending decrease in EGF was observed.

**Conclusions:** Our study suggests that shifts in human T-cell populations and specific immune proteins could serve as promising biomarkers and mechanistic targets for ICI nephrotoxicity. The tumor-bearing HIS-BRGS mouse model reproducibly recapitulates the histopathological and immunological features of human ICI-induced nephrotoxicity and represents a validated preclinical platform for testing novel therapeutic interventions to preserve kidney function during cancer immunotherapy.

**Translational Statement:** Immune checkpoint inhibitor (ICI)-associated nephrotoxicity occurs in up to 25% of treated patients, yet the immunological mechanisms driving renal injury remain poorly characterized due to the scarcity of human biopsy material and the absence of robust preclinical models that recapitulate human immune responses. This study demonstrates that tumor-bearing humanized immune system (HIS) mice treated with combined nivolumab and ipilimumab reproducibly develop renal vasculitis and interstitial nephritis mediated by a human CD4^+^ T cell-dominant infiltrate, mirroring the clinicopathological features reported in patients with ICI-associated acute kidney injury. By integrating histopathology, flow cytometry, multiplex proteomics, and transcriptomics, we identify a coordinated immune network, including IL-15, CCL1, MIF, GZMA, and BAFF, that correlates with the severity of renal pathology and represents tractable mechanistic targets and candidate biomarkers. These findings provide a validated preclinical platform for dissecting irAE mechanisms and testing novel therapeutic strategies to preserve kidney function during cancer immunotherapy.

## Introduction

Immune checkpoint inhibitors (ICIs) targeting programmed cell death protein-1 (PD-1), programmed death-ligand 1 (PD-L1), and cytotoxic T-lymphocyte–associated protein 4 (CTLA-4) have transformed the treatment of cancer by enhancing antitumor responses. However, these drugs can also trigger off-target immune activation leading to immune-related adverse events (irAEs) across multiple organs. Among these, kidney injury, most often presented as acute (tubulo) interstitial nephritis (ATIN/AIN), has emerged as an increasingly recognized complication of ICI therapy affecting ∼5-25% of treated patients. ^1–4^ Histopathological analyses of ICI-associated acute kidney injury (ICI-AKI) cases reveal dense lymphocytic infiltrates in patient biopsies. Recent immunophenotyping of patient kidney biopsies indicate these infiltrates are enriched with activated T cells (CD3+) and a notable expansion of CD4⁺ memory and T-helper cells alongside cytotoxic CD8⁺ T cells, while B-cell, macrophage, and NK-cell involvement remained less frequent compared to controls.^3,5,6^ Transcriptomic, proteomic, and flow cytometric profiling within urine, kidney tissue, and peripheral blood of patients with ICI-AKI identified the upregulation of key immune checkpoints, specifically PD-L1 on tubular cells, alongside pro-inflammatory cytokines such as TNFα and cytotoxic mediators.^5,7,8^ These findings implicate effector T-cell activation and cytokine-driven inflammation as central mechanisms of ICI nephrotoxicity, although mechanistic dissection of tissue-specific immune responses has been limited by the scarcity of human biopsy material and the absence of robust preclinical systems that mirror the human immune system.

Human immune system (HIS) mouse models, generated by engrafting immunodeficient mice with human hematopoietic stem cells, offer a powerful platform for modeling ICI responses including irAEs. ^9–13^ Such models can further recapitulate both antitumor efficacy and immune-related tissue injury of ICI therapies, enabling interrogation of immune, cytokine, and checkpoint signaling networks ^9,14–16^ However, comprehensive characterization of kidney immune responses and biomarker dynamics in these models remains limited.

In this study, we sought to expand on our previously reported kidney pathology in tumor-bearing humanized BRGS (HIS-BRGS) mice^14^ by evaluating kidney-infiltrating T cells, as well as cytokine and chemokine profiles in plasma and kidney tissue, and validating kidney biomarkers.

## Materials and Methods

### Tumor-Bearing Human Immune System Mice

HIS-BRGS (BALB/c-*Rag2^null^Il2rγ^null^Sirpα^NOD^*) mice were generated by the Preclinical Human Immune System Mice Shared Resource (PHISM) at the University of Colorado. Sub-lethally irradiated newborn BRGS pups were injected with human CD34^+^ cells isolated from deidentified human umbilical cord blood obtained from the OB/Gyn Unit at the University of Colorado Hospital (Aurora, CO) following consent of healthy mothers by the Perinatal Research Core (IRB# 16-0541).^9,12,14,17^ Human chimerism at 10-16 weeks of age was determined by assessment of mouse (mCD45^+^) and human hematopoietic cells (hCD45^+^), human T cells (hCD3: hCD4^+^ and hCD8^+^), and human B cells (hCD19^+^) in the blood using flow cytometry. Mice with chimerism (hCD45^+^ divided by total (human+mouse) CD45^+^) over 20% were used for the study. At 17-21 weeks of age, BRGS and HIS-BRGS mice were subcutaneously implanted with MDA-MB-231 (ATCC) cancer cells (3-5 x 10^6^ cells) into both flanks. Tumors were measured with calipers twice weekly, and treatments began when tumor volumes were ∼150 mm^3^. Mice were allocated into vehicle control (PBS) or ICI combination (nivolumab/ipilimumab (20 and 10 mg/kg, i.p. respectively)) groups and treatments were injected 1x/week for 4 weeks, with tumor size and body weights recorded weekly. Mice were euthanized, and plasma, kidneys, lymph nodes, spleens, and tumors collected following treatment. To validate kidney-injury biomarkers, kidneys were collected from adult male C57BL/6 mice (Charles River, Wilmington, MA) treated with vehicle (0.9% saline, n=2) or cisplatin (20 mg/kg, n=2) at 4 days post-treatment.

### Histopathology

Kidneys were fixed in zinc formalin and embedded in paraffin. Sections (5 µm) were stained with hematoxylin and eosin and evaluated for tubular degeneration, protein casts, interstitial nephritis, glomerulonephritis, tubulitis, and vasculitis/periarteritis by a board-certified veterinary anatomic pathologist. Sections were semi-quantitatively scored for interstitial nephritis and vasculitis/periarteritis as previously described.^14^ Images were obtained at 40x magnification using a Jenoptik Gryphax microscope camera (Jenoptik, Fremont, CA).

### Transmission Electron Microscopy (TEM)

Kidneys (1-3 mm^3^) were fixed in 2.5% glutaraldehyde and 4% paraformaldehyde in PBS and embedded before sectioning and then post-fixed in buffered 1% osmium tetroxide. Samples were subsequently dehydrated in a graded series of acetone and embedded in Embed812 resin. Sections (90nm) were cut on a Leica UC6 ultramicrotome and stained with saturated solution of uranyl acetate and lead citrate. Images were captured with an AMT (Advanced Microscopy Techniques) XR111 digital camera at 80Kv on a Philips CM12 transmission electron microscope.

### Flow Cytometry

Single cell suspensions of lymph nodes and spleens were prepared as described.^18^ For tumors and kidneys, approximately half of each excised organ was digested using a Miltenyi GentleMACS Dissociator in C tubes with Liberase DL (Roche) in serum-free media. Digestion was quenched by resuspending the tissue in 10% FCS medium, followed by filtration through a 70 µm filter. After centrifugation, cells were resuspended in 5% FCS media containing DNase, using a volume optimized for maximal flow cytometric recovery. Approximately 1–4 million cells were stained with fluorescently labeled antibodies (**Supplemental Table 1),** either directly *ex vivo* or following a 5-hour stimulation to detect IFN-γ and TNF-α.^13^ Intracellular staining for FoxP3 or Granzyme B was performed using the eBioscience Transcription Factor Kit (**Supplemental Table 1);** for intracellular cytokine detection, cells were fixed in 1% formaldehyde and permeabilized with saponin. Cells from the direct *ex vivo* panel were run on an Aurora Spectral cytometer (Cytek) and *in vitro* cell stimulation panel was assessed using the Penteon (Agilent). Data were analyzed using FlowJo™ software (BD Biosciences).

### Multiplex ELISAs

Multiplex ELISAs were performed using the following kits: 1) human immune response panel 80 plex (ThermoFisher Scientific); 2) mouse kidney injury biomarker panels (KIM-1, IP-10/CXCL10, Renin, B2M, VEGF, TIMP-1, EGF, NGAL, OPN, and Clusterin) (MilliporeSigma); and 3) human immune-oncology checkpoint protein panel (PD-1, CTLA-4, PD-L1, CD80 and CD86) (MilliporeSigma). Frozen kidney tissues (20-30 mg) were homogenized in 100-150 µl of cell extraction buffer using a TissueLyser LT (Qiagen). Samples were then centrifuged at 16000 *xg* for 10 min at 4°C. Multiplex panels were run with plasma and/or diluted supernatant (10 mg/ml in PBS) on a Luminex-200 according to the manufacturer’s protocol, with the limit of detection (LOD) for each protein in **Supplemental Table 2**.

### Quantitative Polymerase Chain Reaction (qPCR) and RNA-sequencing (RNAseq) Analysis

Kidneys were stored frozen in RNA*later* Stabilization Solution (Invitrogen) and then homogenized in RLT buffer (Qiagen) containing 1% of β-mercaptoethanol using TissueLyser LT (Qiagen). Total RNA was isolated using the RNeasy mini kit (Qiagen). Concentrations of total RNA and RNA integrity number (RIN) values were measured using a 2100 Bioanalyzer system. mRNA levels of mouse kidney injury biomarkers and human cytokines/chemokines were quantified by qPCR assay using Sybr Green to detect amplified products (cycle threshold, Ct) in the ABI Vii7 PCR system (Applied Biosystems, Carlesbad, CA) or using bulk RNAseq analysis (Novogene). For RNAseq analysis, gene expression was calculated as fragments per kilobase of transcript per million (FPKM). Differential expression analysis was performed using DESeq2. *P* values were adjusted to control false discovery rate (padj) and statistical significance was set at padj <0.05. For qPCR quantification, Ct values were converted to ΔΔCt by comparing to the mouse reference gene Gapdh. Primers used for qPCR quantification are listed in **Supplemental Table 3**.

### Statistical Analysis

Data are shown individually and expressed as mean ± standard error (SE). Differences across humanized and non-humanized mice and treatment groups were analyzed with two-way ANOVAs followed by Fisher’s LSD posthoc test. Flow cytometry data across four tissues was analyzed using a Brown-Forsythe ANOVA test with an unpaired t-test with Welch’s correction for each tissue. Histopathology data were rank ordered prior to statistical analysis. qPCR data were tested for normal distribution using a Shapiro-Wilk test. Data was then analyzed by an unpaired t-test if normally distributed or a Mann-Whitney test if not normally distributed. Correlations were calculated as Pearson r coefficients and presented as a heatmap (positive correlations are shown in red, and negative correlations are shown in blue). GraphPad Prism version 10 software (GraphPad Software Inc., La Jolla, CA) was used for data analysis. Significance was set at different p-values: *p<0.05, ** p<0.01, ***p<0.001, ****p<0.0001.

## Results

### Animal Characteristics: Weights and Kidney Function

A total of 24 mice completed the study; Veh BRGS n=4; ICI BRGS n=6; Veh HIS-BRGS n=7; and ICI HIS-BRGS n=7. A single immune-related death occurred immediately following the second ICI dose, characterized by high chimerism, diarrhea, and significant weight loss. Weights and kidney function chemistries at study end are provided in **Table 1**. Compared to ICI-treated BRGS mice, the ICI-treated HIS-BRGS mice had statistically lower kidney weights (0.213±0.009 vs. 0.505±0.096 g) and kidney:body weight (1.1±0.03 vs. 2.0±0.3 %), and higher BUN (102.3±10.3 vs 19.3±1.9 mg/dl) and BUN/Cr ratios (361.9±56.5 vs 73.0±8.3).

**Table 1.** Weights and Clinical Chemistries at Study Completion Related to Kidney Function in Vehicle- and ICI-Treated BRGS and HIS-BRGS Mice.^1^.

| Endpoint | Veh BRGS |  | ICI BRGS |  | Veh HIS-BRGS |  | ICI HIS-BRGS |  |
| --- | --- | --- | --- | --- | --- | --- | --- | --- |
|  | Mean | SE | Mean | SE | Mean | SE | Mean | SE |
| **Tumor Weights (g)** | 0.33 | 0.01 | 0.22 | 0.07 | 0.45 | 0.06 | 0.24‡ | 0.02 |
| **Body Weights (g)** | 21.58 | 0.99 | 23.58 | 1.36 | 20.43 | 0.55 | 18.58† | 0.67 |
| **Kidney Weights (g)** | 0.375 | 0.125 | 0.505 | 0.096 | 0.234 | 0.014 | 0.213† | 0.009 |
| **Kidney:Body Weight (%)** | 1.5 | 0.4 | 2.0 | 0.3 | 1.1 | 0.04 | 1.1† | 0.03 |
| **BUN (mg/dl)** | 32.5 | 13.7 | 19.3 | 1.9 | 76.5* | 14.3 | 102.3† | 10.3 |
| **SCr (mg/dl)** | 0.25 | 0.05 | 0.28 | 0.05 | 0.42 | 0.06 | 0.33 | 0.04 |
| **BUN/Cr** | 127.5 | 30.6 | 73.0 | 8.3 | 189.2 | 37.8 | 361.9**,‡ | 56.5 |

<sup>1</sup>Humanized HIS-BRGS and non-humanized BRGS mice were implanted with tumors, followed by treatment with vehicle control (PBS) or immune checkpoint inhibitors (ICI, nivolumab and ipilimumab, 10 mg/kg weekly, i.p.). Clinical chemistries were measured using a Heska Element DC5X. \*p<0.05 compared to vehicle-treated BRGS mice. †p<0.05 compared to ICI-treated BRGS mice. ‡p<0.05 compared to vehicle-treated HIS-BRGS mice.

### Histopathology

Renal vasculitis and interstitial nephritis were observed in ICI-treated HIS-BRGS mice, but not in control BRGS or non-ICI treated HIS-BRGS mice (**Figure 1A,1B).** There were no sex effects on development of vasculitis or interstitial nephritis in HIS-BRGS mice **(Supplemental Figure 1**). While glomerulonephritis was not observed by light microscopy, TEM revealed glomerular basement thickening and fused podocyte foot processes likely due to humanization in HIS-BRGS mice, with more extensive changes following ICI treatment (**Figure 1C**). To inform the effect of humanization on glomerular health, expression of podocyte essential genes^19^ in vehicle-treated BRGS and HIS-BRGS mice was determined by bulk RNA sequencing. Negative values reflected the significantly lower expression of most podocyte genes in HIS-BRGS mice (**Figure 1D**, **Supplemental Table 4**).

**Figure 1.**
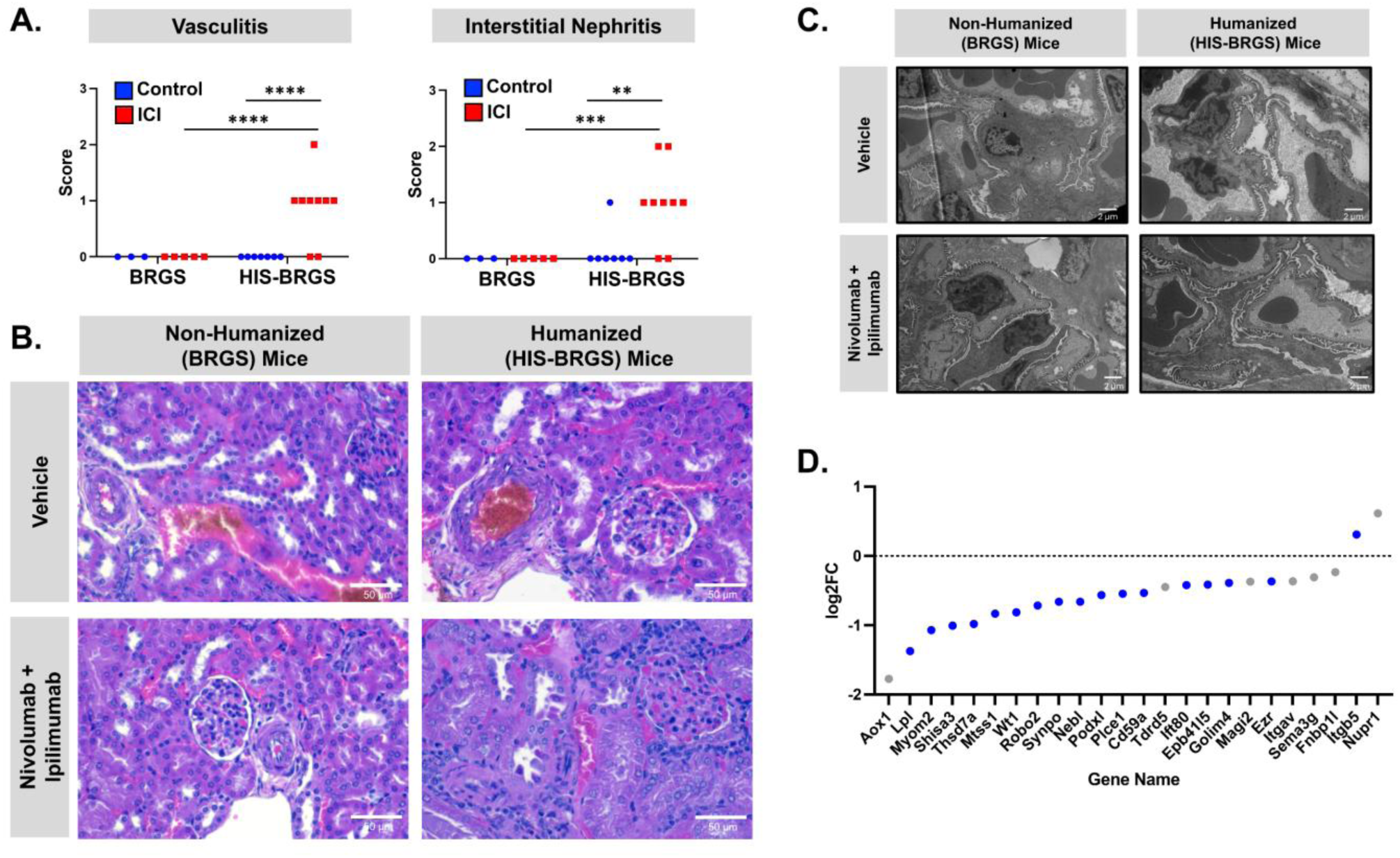
Histopathology and Glomerular Structure Following Treatment with Immune Checkpoint Inhibitors. (A) Humanized HIS-BRGS and non-humanized BRGS mice were implanted with TNBC tumors and treated with vehicle control (PBS) or immune checkpoint inhibitors (ICI, nivolumab and ipilimumab, 20 and 10 mg/kg weekly, i.p.). Kidneys were fixed in formalin and embedded in paraffin before sectioning and staining with H&E. Vasculitis and interstitial nephritis were assessed according to a range of scores (no lesions = 0; minimal lesions, <10% kidney affected = 1; mild lesions, 10-25% kidney affected = 2; moderate lesions, >25-40% kidney affected = 3) by a veterinary anatomic pathologist. ** p<0.01, *** p<0.001, ****p<0.0001. (B) Light microscope images were acquired at 40X magnification. (C) Kidneys (1-3 mm^3^) were fixed in 2.5% glutaraldehyde and 4% paraformaldehyde in PBS and embedded before sectioning and imaging. TEM showed glomerular basement thickening and fused podocyte foot processes following humanization. (D) Podocyte essential genes were quantified in vehicle-treated BRGS and HIS-BRGS mice using bulk RNA sequencing and expressed as log2fold change (FC). Negative values represent lower expression (based on FPKM) in HIS-BRGS mice (blue: adjusted p-value <0.05; grey: not statistically significant). The entire data set is provided in **Supplementary Table 3**.

### Flow Cytometry

Flow cytometry was performed to compare ICI-related changes in human immune cell populations across lymph tissue (lymph nodes and spleen), kidneys, and tumors. Following ICI exposure in HIS-BRGS mice, immune profiling revealed T cell–dominance (97.5%) of total hCD45^+^ cells and decreased CD8^+^ T cells in the kidney, as opposed to decreased CD4^+^ T cells in the spleens and tumors of ICI-treated mice (**Figure 2**). Analysis of activated/immunosuppressed phenotypes of both CD4^+^ and CD8^+^ T cells showed organ specific effects, with more activated T cells (HLA-DR+, effector memory) and fewer exhausted (TIGIT) and Tregs in the tumors and spleens, the tissues with the largest ICI-induced changes, following treatment. Interestingly, exhausted (TIGIT) and regulatory T cells were increased in the lymph nodes, in which significantly fewer terminally exhausted memory (TEMRA) T cells were also measured following treatment. The immune cells in the kidney of ICI-treated mice had similar trends of increased activated immune subsets and decreased exhausted/regulatory cells as observed in the tumors and spleen, with smaller differences. Functionally, ∼46% of kidney localized CD4⁺ T cells produced TNF-α or IFN-γ in ICI treated HIS-BRGS mice, with a significant increase in TNF-α⁺ cells (46%) compared to control HIS-BRGS mice (30.5%) and a trend toward higher dual IFN-γ⁺/TNF-α⁺ expression (31.2% and 18.5%, respectively); while the majority of CD8⁺ T cells produced IFN-γ (54.8%), only 17-21% of CD8⁺ T cells produced dual IFN-γ⁺/TNF-α⁺ or TNF-α⁺ alone and no significant cytokine changes were observed compared to control HIS-BRGS mice (**Figure 3**). Thus, the shift in immune response within the tumor in ICI-treated HIS-BRGS mice is CD8^+^ IFN-γ dominant while in the kidneys, a CD4^+^, TNF-α T cell shift is more dominant.

**Figure 2.**
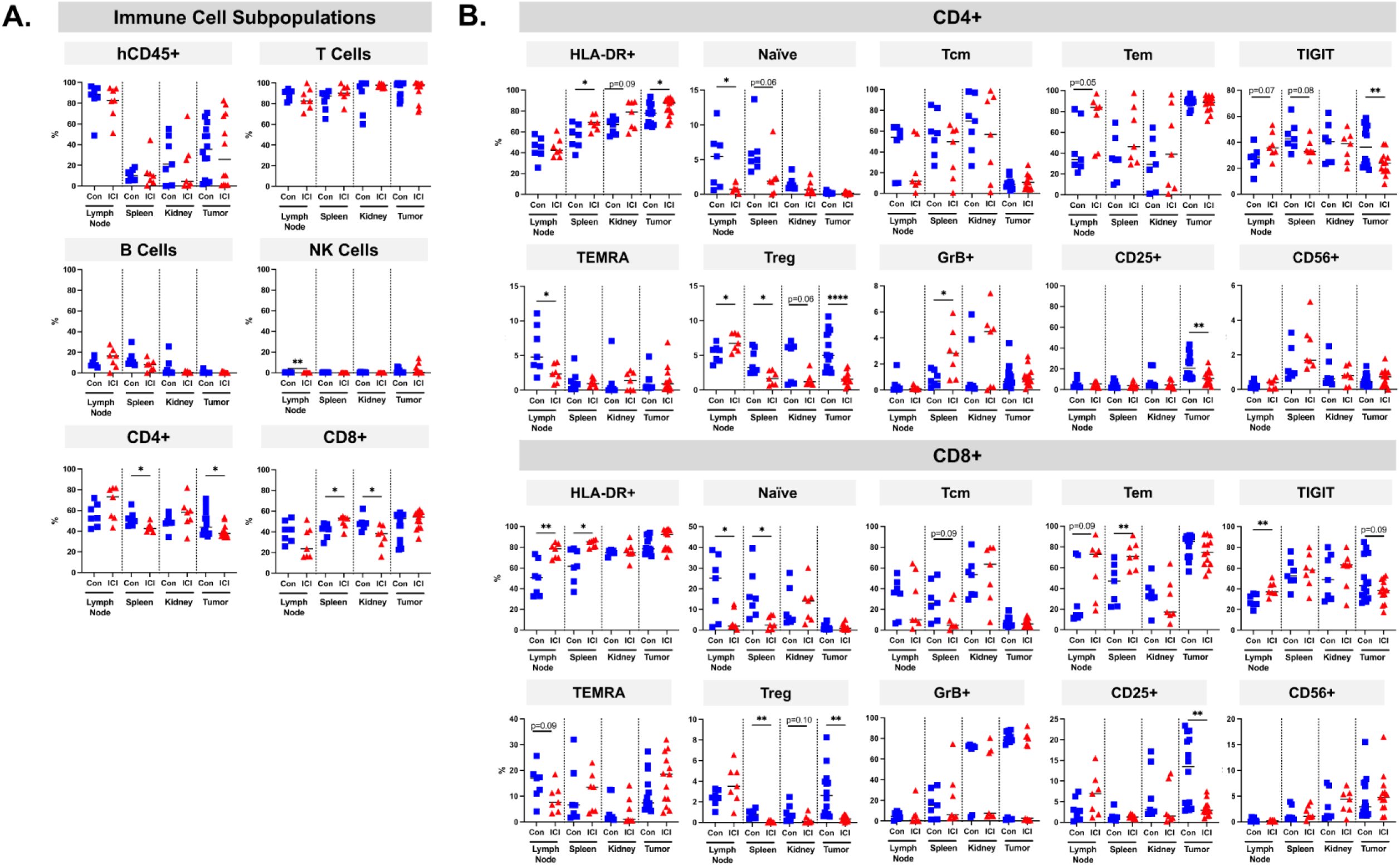
Human Immune Cell Populations in Kidneys of Humanized Mice. Humanized HIS-BRGS mice were implanted with TNBC tumors, followed by treatment with vehicle control (PBS) or immune checkpoint inhibitors (ICI, nivolumab and ipilimumab, 20 and 10 mg/kg weekly, i.p.). Lymph nodes (LN), spleens, tumors, and kidneys were excised and processed into single cell suspensions for flow cytometry. (A) Cells were stained with fluorescent antibodies to quantify human T (both CD4 and CD8), B, and NK among live hCD45^+^ cells in control and ICI-treated HIS-BRGS mice. (B) Cells were stained with fluorescent antibodies to measure activated (HLA-DR, central memory (Tcm), effector memory (Tem), cytotoxic Granzyme B (GrB), CD56) and immunosuppressive (TIGIT, TEMRA, Treg, CD25+FoxP3-) populations among human CD4 and CD8 T cells. *p<0.05, ** p<0.01, ***p<0.001, ****p<0.0001.

**Figure 3.**
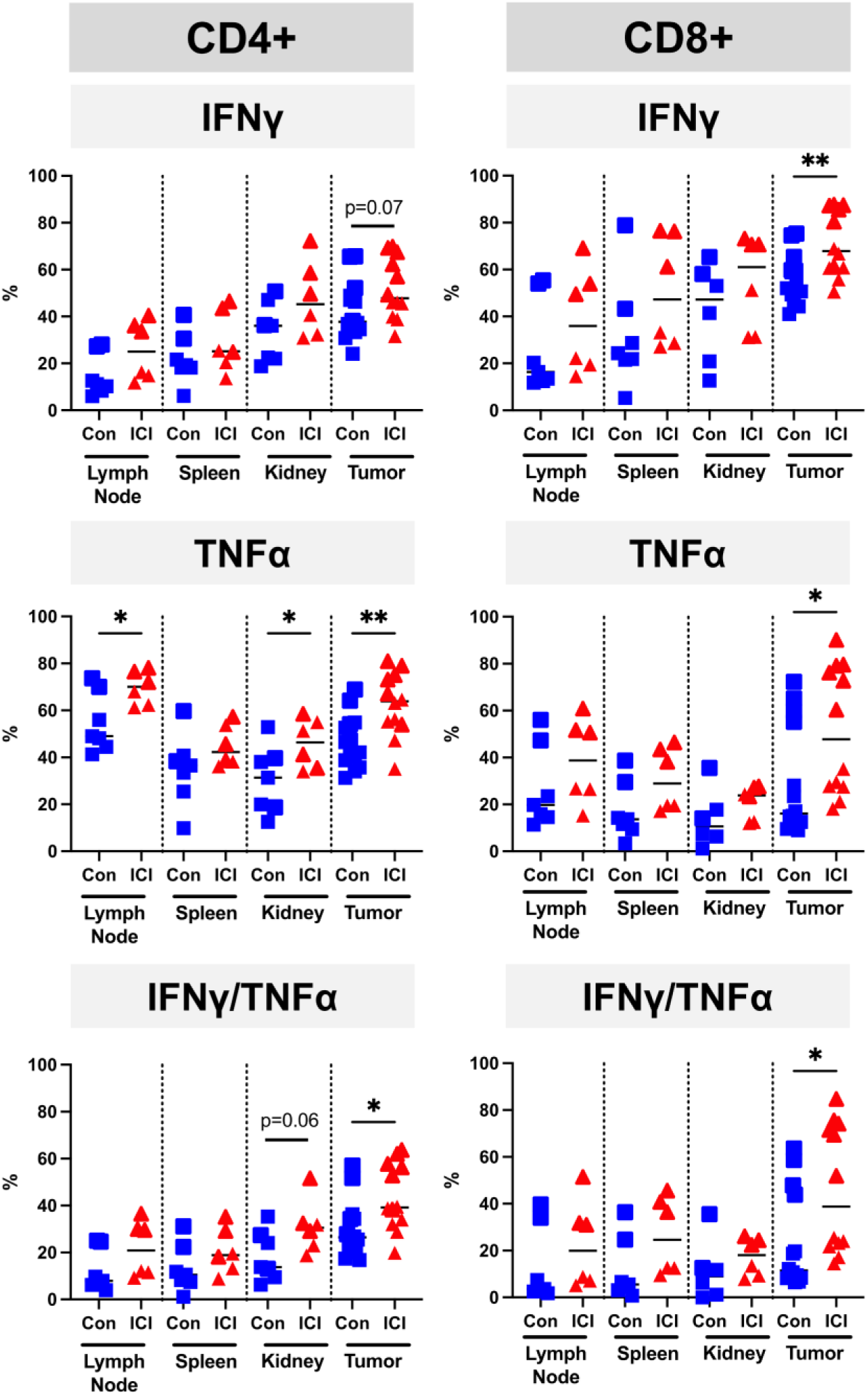
IFNγ and TNFα Production by CD4+ and CD8+ T Cell Populations in Lymph Organs, Tumors and Kidneys of Humanized Mice. Humanized HIS-BRGS mice were implanted with TNBC tumors, followed by treatment with vehicle control (PBS) or immune checkpoint (ICI, nivolumab and ipilimumab, 20 and 10 mg/kg weekly, i.p.). Lymph nodes (LN), spleens, tumors, and kidneys were excised and processed into single cell suspensions. Cell suspensions were stimulated for 5 hours in the presence of a Golgi blocker for the last 4 hours and stained for the presence of intracellular human IFNγ, and TNFα among human CD4 and CD8 T cells. *p<0.05, ** p<0.01, ***p<0.001, ****p<0.0001.

### Immune-related proteins

A human 80 plex immune response panel that measures human cytokines, growth factors, chemokines, proteases, and soluble receptors was evaluated in plasma and kidney homogenates of vehicle- and ICI-treated BRGS and HIS-BRGS mice (**Supplemental Tables 5-6**). Highly significant (p<0.01) relative protein increases in plasma IL-34, LIF, and CCL-25 were observed in vehicle- and ICI-treated HIS-BRGS mice (**Figure 4A**). Other significant (p<0.05) increases were observed in several circulating proteins including IL-15, IL-16, MIF, CCL-1, CCL-7, CCL-17, Galectin-3, BAFF, and CD30 (**Figure 4A**). In kidneys, highly significant (p<0.0001 to p<0.01) relative protein increases in IL-15, CCL-1, MIF, GZMB, BAFF and CD30 were observed in ICI-treated HIS-BRGS mice compared to vehicle-treated counterparts (**Figure 4B**). Significant (p<0.05) increases were also observed in kidney IL-16, CCL-4, GZMA, NGF-β, and TNF-RII with a decrease in IL-4 in ICI-treated mice (**Figure 4B**).

**Figure 4.**
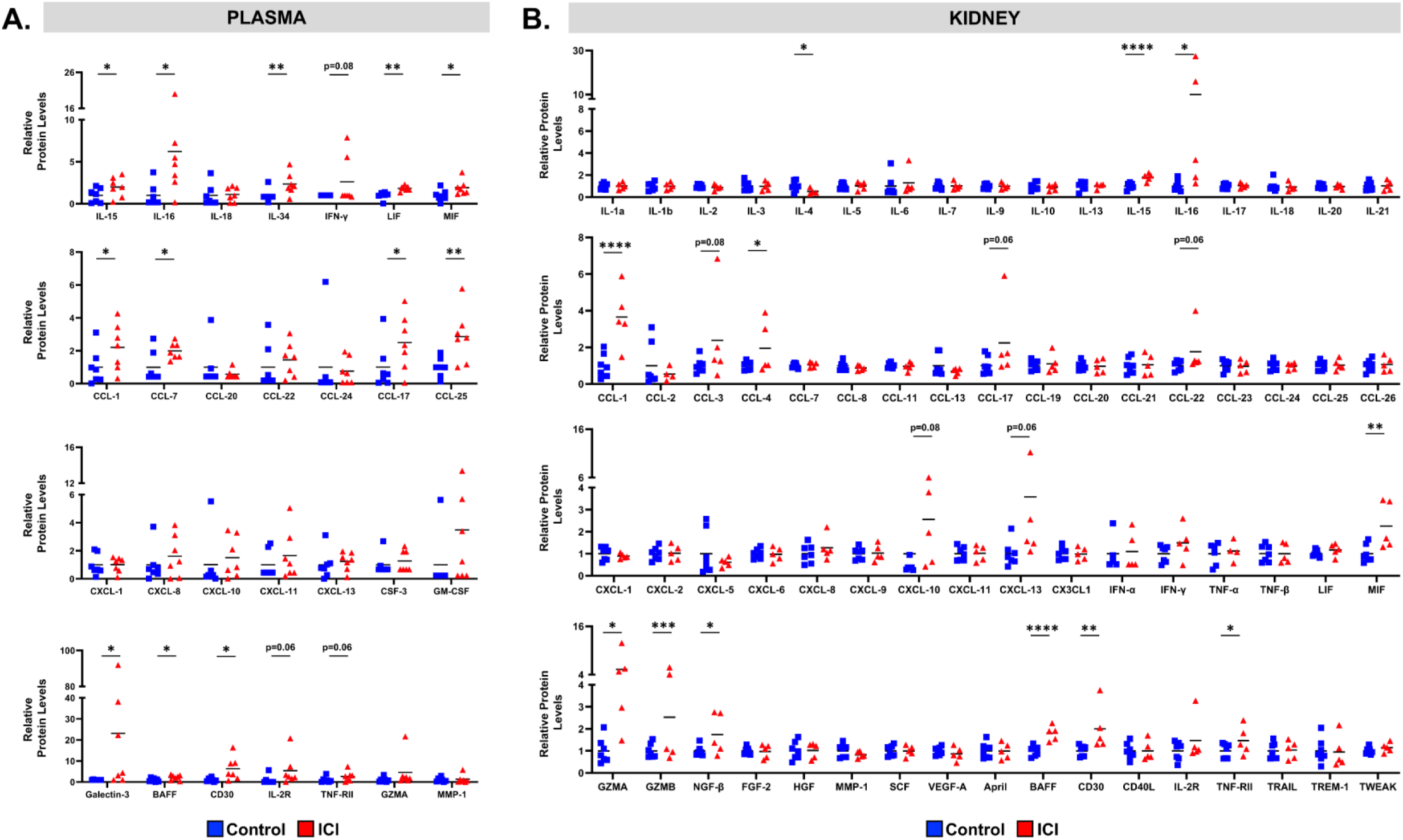
Human Cytokines, Growth Factors, Chemokines, Proteases, and Soluble Receptors in Plasma and Kidneys of HIS-BRGS Mice. Concentrations of immune-related proteins (interleukins and cytokines; chemokine C-C motif ligands; chemokine C-X-C motif ligands, and growth factors; soluble receptors and proteases) were quantified in the (A) plasma and (B) kidney homogenates of vehicle control (PBS) and immune checkpoint inhibitor-treated (ICI, nivolumab and ipilimumab, 20 and 10 mg/kg weekly, i.p.) HIS-BRGS mice using multiplex ELISA against human proteins. Data was adjusted to protein concentration and set to 1 for vehicle-treated HIS-BRGS mice. The entire data set, including concentrations in BRGS mice, is provided in **Supplementary Tables 4-5**. *p<0.05, ** p<0.01, ***p<0.001, ****p<0.0001.

### Kidney Injury Biomarkers

Concentrations of emerging mouse kidney injury biomarker proteins (KIM-1, IP-10/CXCL10, Renin, B2M, VEGF, TIMP-1, EGF, NGAL, OPN, and Clusterin) were quantified in kidney homogenates of BRGS and HIS-BRGS mice following vehicle and ICI treatment (**Supplemental Table 2**). Kidneys from saline- (CON, n=2) and cisplatin-treated (CIS, 20 mg/kg, i.p. 4 days, n=2) C57BL/6 mice were used for comparison. While CIS treatment changed the enrichment of 7 of the 10 proteins, little changes were observed in HIS-BRGS mice except for EGF. Decreases in kidney EGF were observed between vehicle treated BRGS and HIS-BRGS mice and ICI-treated BRGS and HIS-BRGS mice, with a trending decrease (p=0.08) of EGF in kidneys of ICI-treated HIS-BRGS mice compared to vehicle-treated (**Figure 5A**).

**Figure 5.**
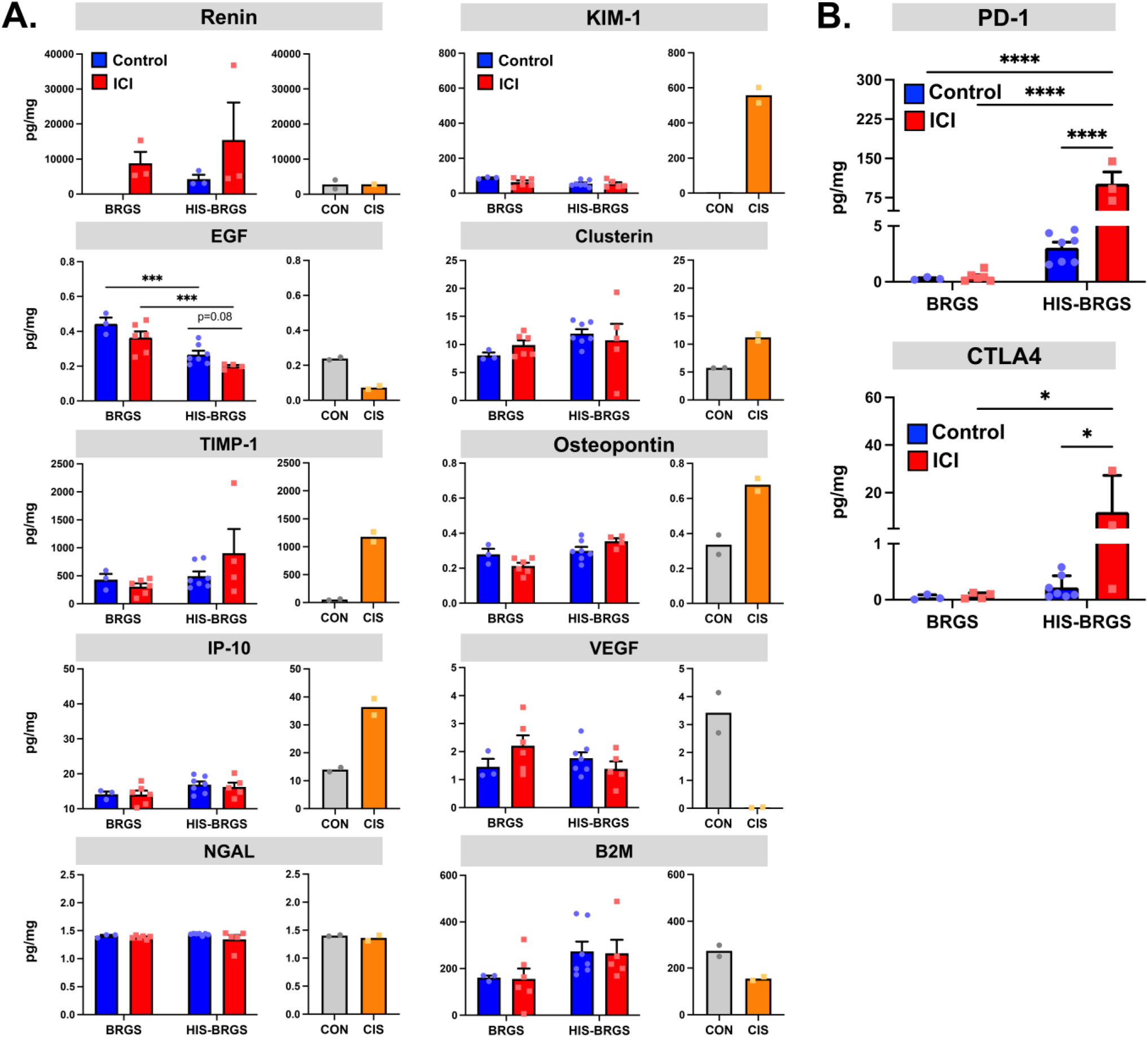
Injury-Related Biomarkers and Human Immune Checkpoint Proteins in Kidneys of BRGS and HIS-BRGS Mice. (A) Concentrations of emerging injury biomarkers were quantified in the homogenates of kidneys from vehicle control (PBS) and immune checkpoint inhibitor-treated (ICI, nivolumab and ipilimumab, 20 and 10 mg/kg weekly, i.p.) BRGS and HIS-BRGS mice using multiplex ELISA against mouse proteins. Controls were obtained from saline-treated (CON, n=2) and cisplatin-treated (CIS, 20 mg/kg, n=2) mice 4 days after injection. Data are presented as protein concentration. ***p<0.001. (B) Concentrations of human immune checkpoint proteins were quantified in the homogenates of kidneys from vehicle control (PBS) and immune checkpoint inhibitor-treated (ICI, nivolumab and ipilimumab, 20 and 10 mg/kg weekly, i.p.) BRGS and HIS-BRGS mice using multiplex ELISA against human proteins. No changes in drug ligands CD80, CD86, or PD-L1 were observed. Data are presented as protein concentration. *p<0.05, ****p<0.0001.

### Checkpoint Proteins

Concentrations of human immune-oncology proteins, CTLA4, PD-1, CD80, CD86, and PD-L1, were quantified in vehicle- and ICI-treated BRGS and HIS-BRGS mice (**Supplemental Table 2**). Highly significant (p<0.0001) protein increases in kidney PD-1 were observed in ICI-treated vs. vehicle-treated HIS-BRGS mice, and ICI-treated HIS-BRGS vs. ICI-treated BRGS mice (**Figure 5B**). Significant (p<0.05) protein increases in kidney CTLA4 were also observed in ICI-treated vs. non-treated HIS-BRGS mice, and ICI-treated HIS-BRGS vs. ICI-treated BRGS mice (**Figure 5B**). No changes in CD80, CD86, or PD-L1 were observed (data not shown).

### Human Enriched Genes in Kidneys of HIS-BRGS Mice

mRNA expression of enriched human and mouse genes in kidneys was quantified in total RNA from vehicle-and ICI-treated HIS-BRGS mice. Bulk RNA sequencing of mouse genes revealed differentially enriched genes (8 increased; 5 decreased) consistent with cellular stress and injury following ICI treatment (**Table 2**). As no significant differences in human transcript expression were detected via RNA sequencing (data not shown), qPCR was subsequently employed to specifically evaluate a targeted panel of human immune-related genes. Highly significant increases (p<0.01) in human ASXL2 and CCS were observed in ICI-treated HIS-BRGS mice (**Figure 6A**). Increases (p<0.05) in human GZMA, ICOS, and LNPEP genes were also observed among others (**Figure 6A**).

**Figure 6.**
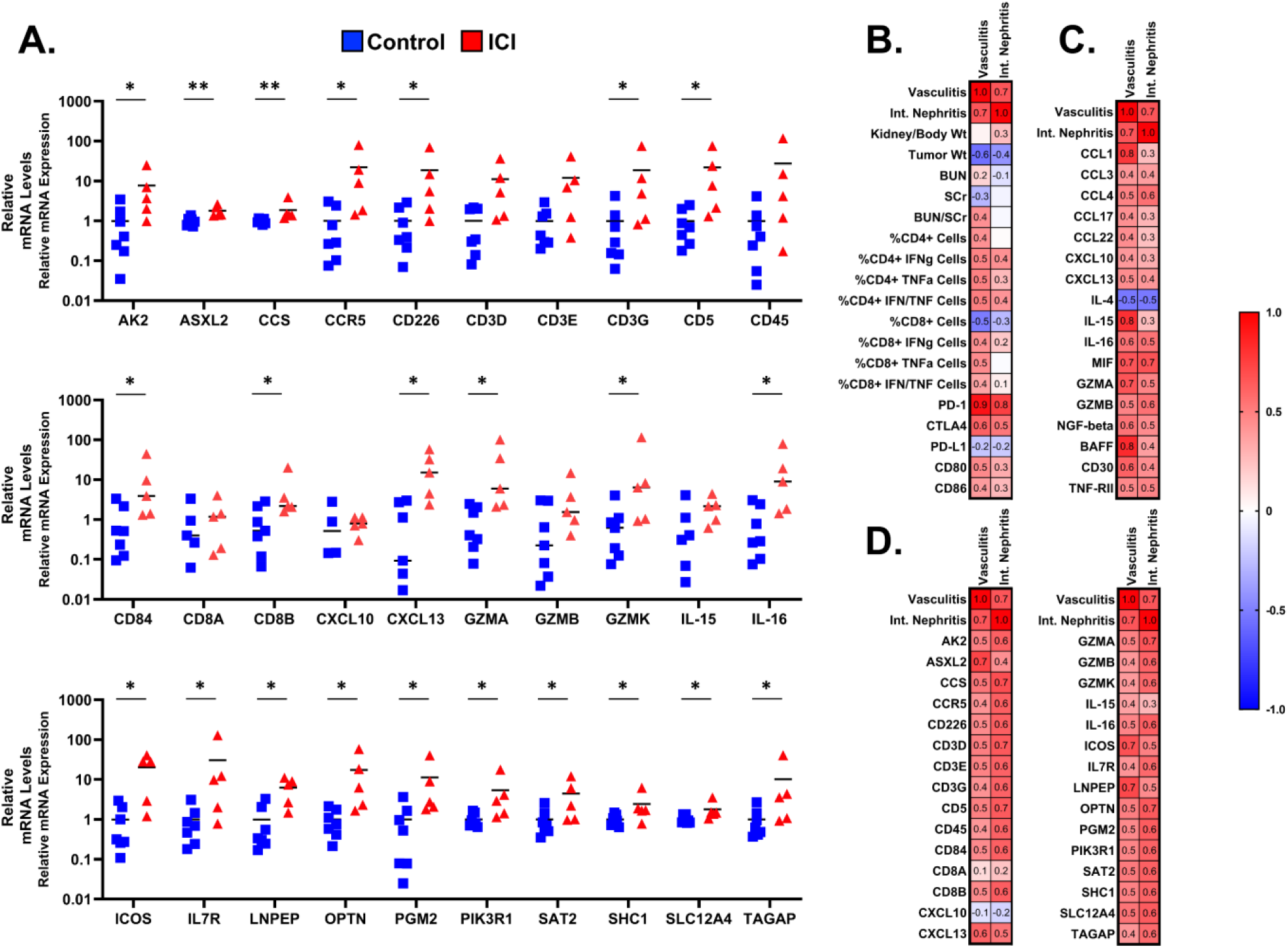
Human Enriched Genes in Kidneys of HIS-BRGS Mice and Correlation Analyses. (A) mRNA expression of enriched genes was quantified in total RNA isolated from kidneys from vehicle control (PBS) and immune checkpoint inhibitor-treated (ICI, nivolumab and ipilimumab, 20 and 10 mg/kg weekly, i.p.) HIS-BRGS mice using qPCR with human primers detailed in **Supplemental Table 3**. Data were normalized to mouse Gapdh and set to 1 for vehicle-treated HIS-BRGS mice. *p<0.05, ** p<0.01. (B-D) Pearson r correlations were calculated between renal histopathology endpoints (vasculitis and interstitial nephritis), and (B) kidney and tumor weights, immune profiles, and checkpoint inhibitor protein levels, (C) renal concentrations of immune mediators and (D) renal mRNA expression in vehicle control (PBS) and immune checkpoint inhibitor-treated (ICI, nivolumab and ipilimumab, 10 mg/kg weekly, ip) HIS-BRGS mice. Immune proteins (pg/mg protein) included in the correlations were those with p<0.1 in the multiplex ELISA quantification.

**Table 2.**
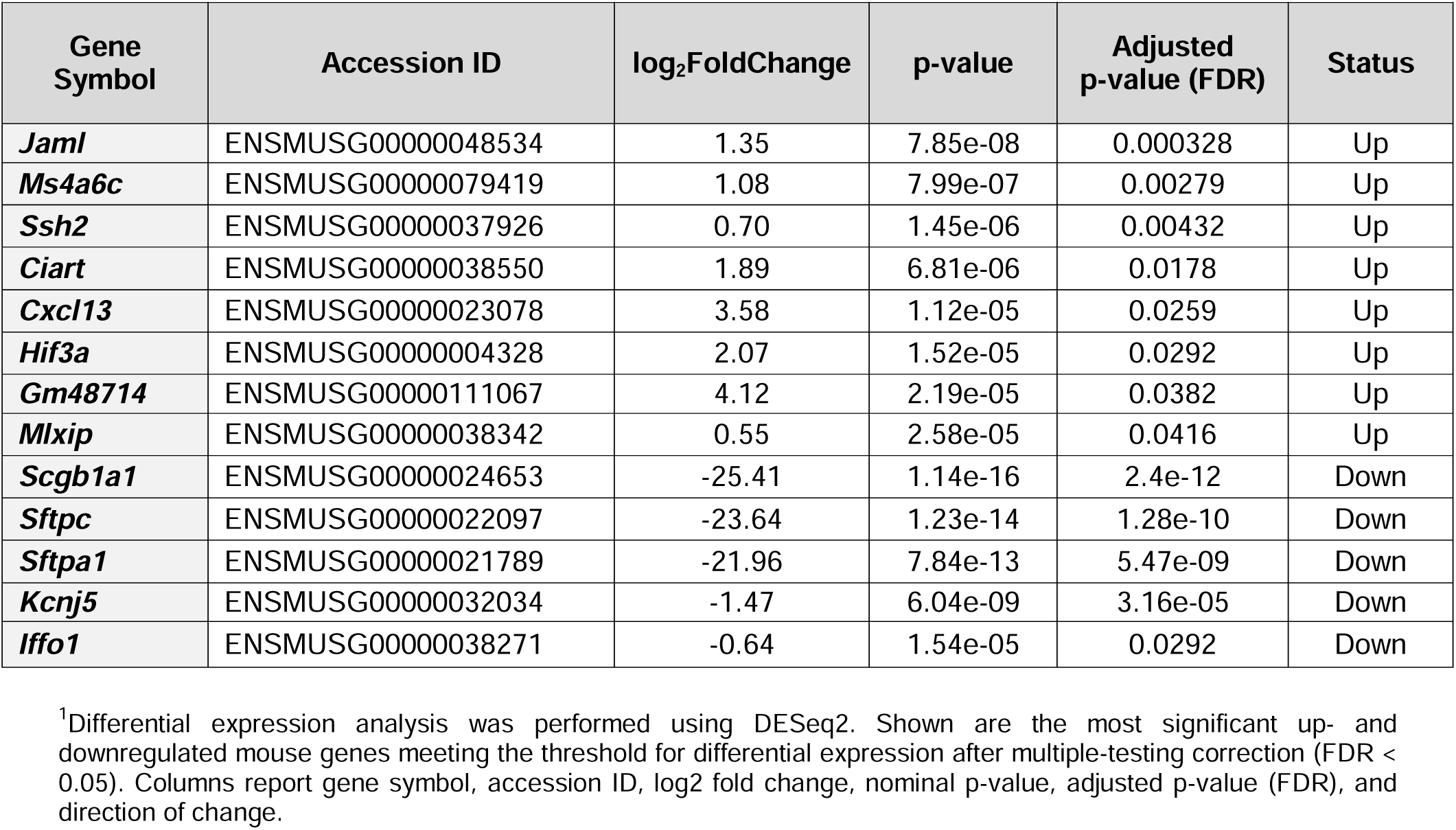
Differentially Expressed Mouse Genes in the Kidneys of ICI-Treated HIS-BRGS Mice. ^1^.

### Predictors of Histopathological Injury

To identify novel indicators that align with renal vasculitis and interstitial nephritis, Pearson r correlations were calculated using data from vehicle- and ICI-treated HIS-BRGS mice including kidney and tumor weights, immune profiles (renal CD4/CD8 percentages), checkpoint proteins, and immune genes and proteins (**Figure 6B-D**). Vasculitis was highly correlated (r = 0.7-1.0, p<0.05) with renal protein concentrations of PD-1, CCL1, IL15, MIF, GZMA, BAFF and renal mRNA expression of ASXL2, ICOS, and LNPEP. Interstitial nephritis was highly correlated (r= 0.7-0.8, p<0.05) with renal concentrations of PD-1 and MIF and renal mRNA expression of CCS, CD3D, CD5, GZMA, and OPTN.

## Discussion

In this study, we extended prior work in the HIS-BRGS model to define novel immune mechanisms underlying kidney injury.^9,12,14,17^ Using combined nivolumab and ipilimumab treatment, we integrated histopathology, flow cytometry, multiplex proteomics, and mouse- and human-specific gene expression to profile the renal immune landscape.

### Histopathology: Reproducibility and Distinctions

Consistent with our previous study,^14^ renal vasculitis and interstitial nephritis were the hallmark pathological features observed in ICI-treated HIS-BRGS mice, reliably recapitulating lesions shown in clinical ICI-associated acute kidney injury (ICI-AKI). ^3^ Our findings of renal vasculitis are particularly noteworthy, as small-vessel vasculitis, although observed less frequently (∼5%), is increasingly recognized as a distinct clinicopathological feature of ICI therapy in humans.^3,20–22^ Although glomerulonephritis was not observed by light microscopy, the glomerular structure of humanized mice under TEM showed glomerular basement thickening and fused podocyte foot processes that were more prominent with ICI treatment. Vehicle treated HIS-BRGS mice demonstrated significant decreases in podocyte essential genes compared to vehicle treated non-humanized BRGS mice, where knockdown of these genes has been previously shown to cause cytoskeletal injury or structural and functional deficiencies.^19,23^ We speculate that the glomerular changes observed in this model are likely a consequence of the humanization process (xenogeneic graft-versus-host activity) and intensified by ICI treatment.

### Elevated Baseline Immune Infiltration and T-cell Dominance

The baseline immune composition of control HIS-BRGS kidneys differed from healthy human renal tissue, with human CD3⁺ T cells comprising ∼88% of intrarenal CD45⁺ leukocytes, almost doubled the ∼47% reported in healthy human kidneys by Park et al., reflecting the preferential engraftment of T and B lymphocytes in this model.^18,24–27^ This elevated baseline is an acknowledged limitation of the HIS platform; however, the model demonstrated high specificity for ICI-induced pathology, as renal vasculitis and the surge in proinflammatory mediators (e.g., CCL1 and MIF) occurred strictly in ICI-treated HIS-BRGS mice and not in untreated humanized controls. Further, the T-cell predominance we observed mirrors the predominance of T-cell–centric infiltrates reported in clinical ICI-AIN biopsies, where CD20⁺ B cells and CD56⁺ NK cells are at lesser proportions or absent, suggesting that while the HIS-BRGS kidney exists in a primed immunological state, it provides a functionally relevant environment to study the transition from homeostatic residency to pathological inflammation. ^3,5,6,28^ The importance of including non-ICI-treated humanized controls to distinguish background chimerism from drug-specific irAEs is therefore highlighted.

### Mechanisms of Injury: CD4^+^ Dominance and Regulatory Failure

Our immunophenotyping suggests that ICI-associated nephritis in the HIS-BRGS model is primarily driven by the CD4^+^ T-cell compartment, a finding that aligns with emerging clinical evidence but contrasts with the CD8-centric mechanisms typically observed in the tumor microenvironment. ^3,29–33^ While the antitumor efficacy of PD-1/CTLA-4 blockade relies on the expansion of CD8^+^ cytotoxic T lymphocytes (CTLs) to induce malignant cell lysis, our model demonstrates that kidney off-target toxicity may operate via a different immunological program. We observed that CD4^+^ T cells were not only the predominant infiltrate (56.7%) but also exhibited a trending increased activation profile (HLA-DR+) and a relatively higher capacity for proinflammatory cytokine production (IFN-γ⁺/TNF-α⁺) than their CD8^+^ counterparts following ICI treatment in HIS mice. This suggests that ICI-induced kidney injury in this model may represent a delayed-type hypersensitivity (DTH)-like reaction orchestrated by CD4^+^ helper effectors rather than a direct CD8^+^ cytolytic attack on tubular cells. This interpretation is supported by recent clinicopathological studies which found CD4^+^ T-cell populations in ICI-AIN cases is a stronger predictor of kidney dysfunction than CD8^+^ T cells.^3,5,34^ The preferential activation of CD4^+^ cells in the kidney may be driven by the local expression of MHC Class II (HLA-DR) on tubular epithelial cells (as well as podocytes and mesangial cells), effectively turning the renal parenchyma into an active antigen-presenting surface.^5,35–40^ In contrast to the CD4^+^ compartment, renal CD8^+^ T-cell frequencies actually declined following ICI treatment, and their cytokine production remained relatively stable. This dissociation suggests that in the context of the kidney, CD8^+^ cells may function more as auxiliary responders rather than primary cytolytic mediators.^3,5,34,41^

A hallmark of irAEs is the interruption of peripheral tolerance. We observed a trending decrease in FoxP3^+^ CD25^+^ regulatory T cells (CD4^+^ and CD8^+^ Tregs) in the kidneys of treated HIS mice. In a healthy human kidney, the T-cell population is relatively lean and balanced by resident regulatory signatures to prevent autoimmunity.^24,42–45^ The trending loss of these suppressive cells in our model indicates a potential state of uncontrolled effector signaling. This regulatory failure hypothesis is supported by findings that PD-1 blockade can specifically impair the suppressive function of Tregs, especially when combined with anti-CTLA4, and in some cases convert them into a pro-inflammatory Th1-like phenotype.^46–48^ The resulting environment, rich in TNF-α but low in regulatory control, directly correlates with the interstitial damage and vasculitis observed in our histopathological analyses.

### Checkpoint Molecule Induction and Tissue Cytokine Profiles

Following ICI exposure, kidneys from HIS-BRGS mice showed marked increases in both PD-1 and CTLA-4 protein concentrations. The multiplex ELISA used to quantify these proteins detects total protein in kidney homogenates and cannot distinguish between cellular and soluble forms of checkpoint molecules that are endogenously expressed by kidney immune and parenchymal cells and residual drug (nivolumab, ipilimumab) bound with soluble antigen retained within the renal tissue; a notable limitation of this approach. Nonetheless, this robust induction likely reflects a state of chronic antigen-driven T-cell activation and the positive feedback loops inherent to checkpoint blockade. The strong correlation between total renal PD-1 levels and the severity of interstitial nephritis suggests that this molecule serves as a quantitative proxy for the scale of the inflammatory infiltrate. Clinically, increases in kidney PD-1 have been linked to various kidney diseases, and significant increases in urinary soluble PD-1 have been found in patients with ICI-AIN.^49^ ^50^ There is a nuance regarding the magnitude of expression of PD-1 observed in ICI-AIN/ATN, specifically with respect to the presence of PD-1+ cells.^3,8,51,52^ While infiltrating T cells express the receptor, positive PD-L1 staining on tubular epithelial cells has been described as a more significant distinguishing factor for ICI-AIN in clinical biopsies.^8^ The discrepancy between the significance of PD-1 expression within this model and lesser levels observed in clinical reports may be a result of the HIS model itself, where T cells have displayed higher levels of checkpoint molecules relative to human T cells.^27^ Temporal differences in biopsy collection may also contribute, as our model may capture a more acute, peak-inflammatory phase, whereas clinical biopsies are often performed after significant damage has occurred.^50^

In renal tissue, there were highly significant increases in IL-15, CCL-1, MIF, GZMB, BAFF, and CD30, along with moderate increases in IL-16, CCL-4, GZMA, NGF-β, and TNF-RII and a decrease in IL-4. These proteins delineate an immune activation and proinflammatory signature and similar to reports in human ICI-nephritis and other irAEs. IL-15 promotes CD8⁺ and memory T-cell proliferation, while CCL-1 and CCL-4 recruit effector cells, supporting sustained interstitial inflammation. ^53–57^ MIF is a pleiotropic cytokine that acts as an upstream driver of mononuclear cell recruitment; its correlation with vasculitis aligns with clinical data suggesting MIF levels correlate with the severity of renal inflammation in patient kidney disease. ^58–61^ Granzyme A (GZMA) and Granzyme B (GZMB) reflect cytotoxic potential, where GZMB has been reported in ICI-AIN biopsy cases. ^62,63^ Increases in BAFF, which regulates B-cell survival, has been linked to B-cell enhancement of antigen-presentation to CD4^+^ T cells and Th1 polarization, correlating with irAE prediction and kidney autoimmune diseases.^64–69^

### Systemic vs. Renal Compartmentalization of ICI Response

The immune response in our HIS-BRGS model displayed a clear compartmentalization. While tumors and lymphoid organs showed robust expansion of effector subsets, the kidney functioned primarily as a site of localized activation and recruitment. This dissociation is clinically critical; it suggests that kidney injury is not a passive spillover of systemic activation, but rather a coordinated tissue-specific event driven by local immune mediators such as MIF and IL-15. These proteins contribute to recruiting and sustaining circulating effector cells within the renal interstitium.

Notably, we did not observe significant elevations in mouse tubular biomarkers KIM-1, NGAL, or B2M, which mirrors the clinical reality of patients with ICI-AIN where AKI markers, including in kidney biopsies, can remain within normal ranges.^5,70^ The highly significant decrease in renal EGF protein in ICI-treated HIS-BRGS mice compared to ICI-treated BRGS mice is particularly noteworthy. EGF is a major growth factor produced by the thick ascending limb of the loop of Henle and distal convoluted tubule; its downregulation is a known hallmark of tubular dedifferentiation and functional impairment.^71^ ^72^ We interpret this sharp decline in EGF as a state of tubular quiescence or dedifferentiation inherent to the chimeric environment. However, the induction of vasculitis and the surge of mediators like MIF only occur upon ICI treatment, and suggests that while the HIS kidney is primed by low EGF-mediated repair capacity, ICI treatment provides the necessary inflammatory activation to manifest overt pathology. In addition to traditional clinical biomarkers, urinary CXCL9 has been validated as a noninvasive diagnostic biomarker for AIN in both general and ICI-treated patient cohorts, but was not elevated in HIS mouse kidneys following ICI treatment. ^54,70,73^ This may reflect the relatively mild degree of AIN pathology in the current preclinical model.

Plasma analysis revealed significant increases in IL-34, LIF, and CCL-25, and moderate increases in IL-15, IL-16, MIF, CCL-1, CCL-7, CCL-17, galectin-3, BAFF, and CD30 in ICI-treated mice. IL-34 and LIF are emerging cytokines associated with myeloid regulation and tissue repair; their rise in circulation suggests systemic immune activation rather than a role in direct tubular injury.^74,75^ The overlap between plasma and kidney mediators (IL-15, MIF, BAFF, CCL-1), however, supports a coordinated systemic and renal immune response. Monitoring early surges in T-cell–recruiting proteins in parallel with loss of reparative factors like EGF could provide an earlier window for therapeutic intervention than damage markers alone.^5,7^

### Enriched Human Genes in Kidneys

To complement the proteomic landscape, we interrogated the human-specific transcriptional profile. GZMA transcripts were increased in ICI-treated kidneys. This transcriptional surge is concordant with elevated serine protease levels and supports the presence of cytotoxic programming in the infiltrating human lymphocytes. Although flow cytometry indicated a CD4-predominant infiltrate, GZMA upregulation suggests these CD4 cells, and/or the smaller CD8 subset, are equipped for cytolytic function. Clinically, intrarenal granzyme expression has been associated with more aggressive T cell–mediated injury.^76–79^ We also noted significant upregulation of CCS (copper chaperone for superoxide dismutase), consistent with an enhanced antioxidant response to superoxide and other reactive oxygen species generated in the inflamed interstitium.^80–83^ Concomitant induction of ASXL2 suggests the renal parenchyma is engaging epigenetic remodeling programs.^84,85^

### Predictors of Histopathological Changes

The correlation analyses reveal a tightly coordinated immune network in which PD-1 pathway activation aligns closely with cytokine signaling and distinct kidney injury patterns.^8,86^ In our HIS-BRGS model, PD-1 expression on human T cells correlated strongly with both vasculitis and interstitial nephritis, as well as with local cytokine signatures mirroring clinical observations. ^49,87^ In patients with ICI-AKI, the extent of interstitial PD-1⁺ T-cell infiltration correlates with the severity of kidney dysfunction.^3,8,51,52,88^ These results support using the burden of PD-1 as a quantitative indicator of local immune injury risk.^89–91^ Correlation analyses also reveal a specialized cytokine network; IL-15, CCL1, MIF, GZMA, and BAFF, strongly associated with renal vasculitis. IL-15 correlated robustly with CCL1 and BAFF (data not shown), positioning it as a potential upstream coordinator of lymphocyte recruitment and cytotoxic function.^92–95^ Under PD-1 blockade, homeostatic cytokines such as IL-15 promote pathological cytotoxic lymphocyte activation, enabling granzyme-mediated endothelial injury characteristic of vasculitis.^96,97^ CCL1 recruits CCR8+ T cells to perivascular sites, while BAFF sustains B-cell survival and differentiation. ^67,98^ By defining these interlocking correlations, the present work moves beyond single-analyte biomarkers toward a systems-level view of ICI-AKI pathogenesis.

## Conclusions

This study validates the tumor-bearing HIS-BRGS mouse as a robust preclinical platform that successfully recapitulates the histopathological and immunological features of human ICI-induced nephrotoxicity. We demonstrated that treatment with nivolumab and ipilimumab induces reproducible renal vasculitis and interstitial nephritis mediated by a human T-cell dominant infiltrate, characterized by activated CD4^+^ effector cells and a proinflammatory cytokine milieu. Emerging kidney injury biomarkers, such as kidney injury molecule-1, were insensitive in this model. However, intrarenal and circulating immune mediators, specifically IL-15, CCL1, MIF, GZMA, and BAFF, showed strong correlations with pathology, suggesting their potential as mechanism-based biomarkers. Up-regulation of genes, such as GZMA and CCS, highlighted the interplay between cytotoxicity and oxidative stress in ICI-mediated tissue injury. Collectively, these data suggest that monitoring specific shifts in T-cell phenotypes and soluble immune mediators in this novel model may offer a superior strategy for dissecting the complex mechanisms of irAEs and testing novel therapeutic interventions to preserve kidney function during cancer immunotherapy.

## Supporting information

ARRIVE

## Data Statement

ARRIVE reporting guidelines were used in drafting this manuscript, and the ARRIVE reporting checklist when editing, included in Supplemental File. ^99^ The images used for analysis in this work are publicly available in FigShare at 10.6084/m9.figshare.32145847.

## Acknowledgments

This work was supported by the National Cancer Institute [R01CA277313], National Center for Advancing Translational Sciences [UM1TR004789], National Institute of General Medical Sciences [K12GM093854], and the National Institute of Environmental Health Sciences [P30ES005022, T32ES001748, and T32ES029074], components of the National Institutes of Health (NIH), and HESI Thrive. All animal studies were conducted at the University of Colorado Animal Care Facilities. The Pre-clinical Human Immune System Mice Shared Resource (PHISM) at the University of Colorado generated and maintained the humanized mouse model and performed flow cytometry and analysis. Luminex instrument services, results, and products supporting the research project were generated by the Rutgers Cancer Institute, Immune Monitoring, and Flow Cytometry Shared Resource, with funding from the NCI-CCSG P30CA072770-5920. Transmission electron microscopy was performed at the Core Imaging Lab at Robert Wood Johnson Medical School. The content is solely the responsibility of the authors and does not necessarily represent the official views of the National Institutes of Health.

## Abbreviations

AIN: acute interstitial nephritis
AK2: Adenylate Kinase 2
AKI: acute kidney injury
APRIL: A PRoliferation-Inducing Ligand (TNF Superfamily Member 13)
ASXL2: ASXL Transcriptional Regulator 2
ATIN: acute tubulointerstitial nephritis
B2M: Beta-2-Microglobulin
BAFF: TNF Superfamily Member 13B
BUN: Blood Urea Nitrogen
CCL1: C-C Motif Chemokine Ligand 1
CCL3: C-C Motif Chemokine Ligand 3
CCL4: C-C Motif Chemokine Ligand 4
CCL17: C-C Motif Chemokine Ligand 17
CCL22: C-C Motif Chemokine Ligand 22
CCS: Copper Chaperone for Superoxide Dismutase
CCR5: C-C Motif Chemokine Receptor 5
CD226: CD226 Molecule
CD3D: CD3 Delta Subunit Of T-Cell Receptor Complex
CD3E: CD3 Epsilon Subunit Of T-Cell Receptor Complex
CD3G: CD3 Gamma Subunit Of T-Cell Receptor Complex
CD5: CD5 Molecule
CD8A: CD8 Subunit Alpha
CD8B: CD8 Subunit Beta
CD30: TNF Receptor Superfamily Member 8
CD40L: CD40 Ligand
CD45: protein tyrosine phosphatase receptor type C (PTPRC)
CD80: CD80 Molecule
CD84: CD84 Molecule
CD86: CD86 Molecule
CTLA4: Cytotoxic T-Lymphocyte Associated Protein 4
CXCL10 (IP-10): C-X-C Motif Chemokine Ligand 10
CXCL13: C-X-C Motif Chemokine Ligand 13
EGF: Epidermal Growth Factor
FGF-2: Fibroblast Growth Factor 2
FPKM: Fragments Per Kilobase of transcript per Million
GM-CSF: Colony Stimulating Factor 2
GZMA: Granzyme A
GZMB: Granzyme B
GZMK: Granzyme K
HGF: Hepatocyte Growth Factor
HIS: human immune system
ICI: immune checkpoint inhibitor
IL-2RA: Interleukin 2 Receptor Subunit Alpha
IL-4: Interleukin 4
IL-15: Interleukin 15
IL-16: Interleukin 16
ICOS: Inducible T Cell Costimulator
IL7R: Interleukin 7 Receptor
IFN-α: Interferon Alpha
IFN-γ: Interferon Gamma
irAE: immune-related adverse events
KIM-1: Hepatitis A Virus Cellular Receptor 1
LIF: LIF Interleukin 6 Family Cytokine
LNPEP: Leucyl And Cystinyl Aminopeptidase
MIF: Macrophage Migration Inhibitory Factor
MMP1: Matrix Metallopeptidase 1
NGAL/LCN2: Lipocalin 2
OPTN: Optineurin
PD-1: Programmed Cell Death 1
PGM2: Phosphoglucomutase 2
PIK3R1: Phosphoinositide-3-Kinase Regulatory Subunit 1
SAT2: Spermidine/Spermine N1-Acetyltransferase Family Member 2
SCF: KIT Ligand
SCr: Serum creatinine
SHC1: SHC Adaptor Protein 1
SLC12A4: Solute Carrier Family 12 Member 4
TAGAP: T Cell Activation RhoGTPase Activating Protein
TIMP1: TIMP Metallopeptidase Inhibitor 1
TNF-α: Tumor Necrosis Factor Alpha
TNF-β: Tumor Necrosis Factor Beta
TRAIL: TNF Superfamily Member 10
TNF-RII: TNF Receptor Superfamily Member 1B
TREM-1: Triggering Receptor Expressed On Myeloid Cells 1
TWEAK: TNF Superfamily Member 12
VEGF-A: Vascular Endothelial Growth Factor Alpha

## References

1 Cortazar, F. B. et al. Clinicopathological features of acute kidney injury associated with immune checkpoint inhibitors. Kidney Int 90, 638–647 (2016). 10.1016/j.kint.2016.04.008

2 Gupta, S. et al. Acute kidney injury in patients treated with immune checkpoint inhibitors. J Immunother Cancer 9 (2021). 10.1136/jitc-2021-003467

3 Xu, L. Y. et al. Clinicopathological Features of Kidney Injury Related to Immune Checkpoint Inhibitors: A Systematic Review. J Clin Med 12 (2023). 10.3390/jcm12041349

4 Baker, M. L. et al. Mortality after acute kidney injury and acute interstitial nephritis in patients prescribed immune checkpoint inhibitor therapy. Journal for ImmunoTherapy of Cancer 10, e004421 (2022). 10.1136/jitc-2021-004421

5 Farooqui, N. et al. Cytokines and Immune Cell Phenotype in Acute Kidney Injury Associated With Immune Checkpoint Inhibitors. Kidney Int Rep 8, 628–641 (2023). 10.1016/j.ekir.2022.11.020

6 Tominaga, K. et al. Predominant CD8(+) cell infiltration and low accumulation of regulatory T cells in immune checkpoint inhibitor-induced tubulointerstitial nephritis. Pathol Int 74, 317–326 (2024). 10.1111/pin.13428

7 Sise, M. E. et al. Soluble and cell-based markers of immune checkpoint inhibitor-associated nephritis. J Immunother Cancer 11 (2023). 10.1136/jitc-2022-006222

8 Cassol, C. et al. Anti-PD-1 Immunotherapy May Induce Interstitial Nephritis With Increased Tubular Epithelial Expression of PD-L1. Kidney Int Rep 4, 1152–1160 (2019). 10.1016/j.ekir.2019.06.001

9 Capasso, A. et al. Characterization of immune responses to anti-PD-1 mono and combination immunotherapy in hematopoietic humanized mice implanted with tumor xenografts. J Immunother Cancer 7, 37 (2019). 10.1186/s40425-019-0518-z

10 Chuprin, J. et al. Humanized mouse models for immuno-oncology research. Nat Rev Clin Oncol 20, 192–206 (2023). 10.1038/s41571-022-00721-2

11 De La Rochere, P., Guil-Luna, S., Decaudin, D., Azar, G., Sidhu, S. S. & Piaggio, E. Humanized Mice for the Study of Immuno-Oncology. Trends in Immunology 39, 748–763 (2018). 10.1016/j.it.2018.07.001

12 Lang, J. et al. Development of an Adrenocortical Cancer Humanized Mouse Model to Characterize Anti-PD1 Effects on Tumor Microenvironment. The Journal of Clinical Endocrinology & Metabolism 105, 26–42 (2019). 10.1210/clinem/dgz014

13 Lanis, J. M. et al. Testing Cancer Immunotherapeutics in a Humanized Mouse Model Bearing Human Tumors. J Vis Exp (2022). 10.3791/64606

14 Asby, S. C. et al. Pathological findings of immunotherapy-induced nephrotoxicity in a humanized immune system mouse model. Kidney Int 107, 930–934 (2025). 10.1016/j.kint.2025.01.021

15 Park, N. et al. Preclinical platform for long-term evaluation of immuno-oncology drugs using hCD34+ humanized mouse model. J Immunother Cancer 8 (2020). 10.1136/jitc-2020-001513

16 Vudattu, N. K. et al. Humanized mice as a model for aberrant responses in human T cell immunotherapy. J Immunol 193, 587–596 (2014). 10.4049/jimmunol.1302455

17 Lang, J., Weiss, N., Freed, B. M., Torres, R. M. & Pelanda, R. Generation of hematopoietic humanized mice in the newborn BALB/c-Rag2null Il2rγnull mouse model: a multivariable optimization approach. Clin Immunol 140, 102–116 (2011). 10.1016/j.clim.2011.04.002

18 Marín-Jiménez, J. A. et al. Testing Cancer Immunotherapy in a Human Immune System Mouse Model: Correlating Treatment Responses to Human Chimerism, Therapeutic Variables and Immune Cell Phenotypes. Front Immunol 12, 607282 (2021). 10.3389/fimmu.2021.607282

19 Lu, Y. et al. Genome-wide identification of genes essential for podocyte cytoskeletons based on single-cell RNA sequencing. Kidney International 92, 1119–1129 (2017). 10.1016/j.kint.2017.04.022

20 Gallan, A. J., Alexander, E., Reid, P., Kutuby, F., Chang, A. & Henriksen, K. J. Renal Vasculitis and Pauci-immune Glomerulonephritis Associated With Immune Checkpoint Inhibitors. Am J Kidney Dis 74, 853–856 (2019). 10.1053/j.ajkd.2019.04.016

21 Kitchlu, A. et al. A Systematic Review of Immune Checkpoint Inhibitor-Associated Glomerular Disease. Kidney Int Rep 6, 66–77 (2021). 10.1016/j.ekir.2020.10.002

22 Lee, C. M., Wang, M., Rajkumar, A., Calabrese, C. & Calabrese, L. A scoping review of vasculitis as an immune-related adverse event from checkpoint inhibitor therapy of cancer: Unraveling the complexities at the intersection of immunology and vascular pathology. Semin Arthritis Rheum 66, 152440 (2024). 10.1016/j.semarthrit.2024.152440

23 Löwik, M. M., Groenen, P. J., Levtchenko, E. N., Monnens, L. A. & van den Heuvel, L. P. Molecular genetic analysis of podocyte genes in focal segmental glomerulosclerosis--a review. Eur J Pediatr 168, 1291–1304 (2009). 10.1007/s00431-009-1017-x

24 Park, J.-G. et al. Immune cell composition in normal human kidneys. Scientific Reports 10, 15678 (2020). 10.1038/s41598-020-72821-x

25 Lepus, C. M. et al. Comparison of human fetal liver, umbilical cord blood, and adult blood hematopoietic stem cell engraftment in NOD-scid/gammac-/-, Balb/c-Rag1-/-gammac-/-, and C.B-17-scid/bg immunodeficient mice. Hum Immunol 70, 790–802 (2009). 10.1016/j.humimm.2009.06.005

26 Mestas, J. & Hughes, C. C. Of mice and not men: differences between mouse and human immunology. J Immunol 172, 2731–2738 (2004). 10.4049/jimmunol.172.5.2731

27 Patel, A. K. et al. Spontaneous tumor regression mediated by human T cells in a humanized immune system mouse model. Communications Biology 6, 444 (2023). 10.1038/s42003-023-04824-z

28 Abudayyeh, A., Suo, L., Lin, H., Mamlouk, O., Abdel-Wahab, N. & Tchakarov, A. Pathologic Predictors of Response to Treatment of Immune Checkpoint Inhibitor-Induced Kidney Injury. Cancers (Basel) 14 (2022). 10.3390/cancers14215267

29 Aliazis, K. et al. The tumor microenvironment’s role in the response to immune checkpoint blockade. Nat Cancer 6, 924–937 (2025). 10.1038/s43018-025-00986-3

30 Lin, Y., Song, Y., Zhang, Y., Li, X., Kan, L. & Han, S. New insights on anti-tumor immunity of CD8(+) T cells: cancer stem cells, tumor immune microenvironment and immunotherapy. J Transl Med 23, 341 (2025). 10.1186/s12967-025-06291-y

31 Marco, T. et al. The mechanisms of acute interstitial nephritis in the era of immune checkpoint inhibitors in melanoma. Ther Adv Med Oncol 11, 1758835919875549 (2019). 10.1177/1758835919875549

32 Seethapathy, H., Mistry, K. & Sise, M. E. Immunological mechanisms underlying clinical phenotypes and noninvasive diagnosis of immune checkpoint inhibitor-induced kidney disease. Immunol Rev 318, 61–69 (2023). 10.1111/imr.13243

33 Wei, S. C. et al. Distinct Cellular Mechanisms Underlie Anti-CTLA-4 and Anti-PD-1 Checkpoint Blockade. Cell 170, 1120–1133.e1117 (2017). 10.1016/j.cell.2017.07.024

34 Belliere, J. et al. Acute interstitial nephritis related to immune checkpoint inhibitors. Br J Cancer 115, 1457–1461 (2016). 10.1038/bjc.2016.358

35 Cheng, H. F., Nolasco, F., Cameron, J. S., Hildreth, G., Neild, G. H. & Hartley, B. HLA-DR display by renal tubular epithelium and phenotype of infiltrate in interstitial nephritis. Nephrol Dial Transplant 4, 205–215 (1989). 10.1093/oxfordjournals.ndt.a091857

36 Curran, C. S. & Kopp, J. B. PD-1 immunobiology in glomerulonephritis and renal cell carcinoma. BMC Nephrol 22, 80 (2021). 10.1186/s12882-021-02257-6

37 Hasakioğulları, İ., Kockx, M., Szabados, B., Young, M. & Powles, T. Tumor-specific MHC class II upregulation associated with response to anti-PD-L1 therapy in patients with urothelial cancer. Journal of Clinical Oncology 42, 4584–4584 (2024). 10.1200/JCO.2024.42.16_suppl.4584

38 Johnson, D. B. et al. Melanoma-specific MHC-II expression represents a tumour-autonomous phenotype and predicts response to anti-PD-1/PD-L1 therapy. Nature Communications 7, 10582 (2016). 10.1038/ncomms10582

39 Wuthrich, R. P., Glimcher, L. H., Yui, M. A., Jevnikar, A. M., Dumas, S. E. & Kelley, V. E. MHC class II, antigen presentation and tumor necrosis factor in renal tubular epithelial cells. Kidney Int 37, 783–792 (1990). 10.1038/ki.1990.46

40 Yokoyama, H. et al. Up-regulated MHC-class II expression and gamma-IFN and soluble IL-2R in lupus nephritis. Kidney Int 42, 755–763 (1992). 10.1038/ki.1992.344

41 Sade-Feldman, M. et al. Defining T Cell States Associated with Response to Checkpoint Immunotherapy in Melanoma. Cell 175, 998–1013.e1020 (2018). 10.1016/j.cell.2018.10.038

42 Lake, B. B. et al. An atlas of healthy and injured cell states and niches in the human kidney. Nature 619, 585–594 (2023). 10.1038/s41586-023-05769-3

43 Stewart, B. J. et al. Spatiotemporal immune zonation of the human kidney. Science 365, 1461–1466 (2019). doi:10.1126/science.aat5031

44 Valencia, X. & Lipsky, P. E. CD4+CD25+FoxP3+ regulatory T cells in autoimmune diseases. Nature Clinical Practice Rheumatology 3, 619–626 (2007). 10.1038/ncprheum0624

45 Wang, L. et al. Regulatory T cells in homeostasis and disease: molecular mechanisms and therapeutic potential. Signal Transduction and Targeted Therapy 10, 345 (2025). 10.1038/s41392-025-02326-4

46 Attias, M., Al-Aubodah, T. & Piccirillo, C. A. Mechanisms of human FoxP3+ Treg cell development and function in health and disease. Clinical and Experimental Immunology 197, 36–51 (2019). 10.1111/cei.13290

47 Attias, M. et al. Anti-PD-1 amplifies costimulation in melanoma-infiltrating T(h)1-like Foxp3(+) regulatory T cells to alleviate local immunosuppression. J Immunother Cancer 13 (2025). 10.1136/jitc-2024-009435

48 Zhang, P., Wang, Y., Miao, Q. & Chen, Y. The therapeutic potential of PD-1/PD-L1 pathway on immune-related diseases: Based on the innate and adaptive immune components. Biomedicine & Pharmacotherapy 167, 115569 (2023). 10.1016/j.biopha.2023.115569

49 Chen, R. et al. PD-1 immunology in the kidneys: a growing relationship. Front Immunol 15, 1458209 (2024). 10.3389/fimmu.2024.1458209

50 Gomez-Preciado, F. et al. Urinary soluble PD-1 as a biomarker of checkpoint inhibitor-induced acute tubulointerstitial nephritis. Clinical Kidney Journal 17 (2024). 10.1093/ckj/sfae200

51 Casals, J. et al. Differentiating Acute Interstitial Nephritis From Immune Checkpoint Inhibitors From Other Causes. Kidney International Reports 8, 672–675 (2023). 10.1016/j.ekir.2022.12.017

52 Tampe, D., Kopp, S. B., Baier, E., Hakroush, S. & Tampe, B. Compartmentalization of Intrarenal Programmed Cell Death Protein 1-Ligand 1 and Its Receptor in Kidney Injury Related to Immune Checkpoint Inhibitor Nephrotoxicity. Front Med (Lausanne) 9, 902256 (2022). 10.3389/fmed.2022.902256

53 Charo, I. F. & Ransohoff, R. M. The Many Roles of Chemokines and Chemokine Receptors in Inflammation. New England Journal of Medicine 354, 610–621 (2006). doi:10.1056/NEJMra052723

54 Long, J. P., Singh, S., Dong, Y., Yee, C. & Lin, J. S. Urine proteomics defines an immune checkpoint-associated nephritis signature. J Immunother Cancer 13 (2025). 10.1136/jitc-2024-010680

55 Les, I. et al. Predictive Biomarkers for Checkpoint Inhibitor Immune-Related Adverse Events. Cancers (Basel) 15 (2023). 10.3390/cancers15051629

56 Reschke, R., Sullivan, R. J., Lipson, E. J., Enk, A. H., Gajewski, T. F. & Hassel, J. C. Targeting molecular pathways to control immune checkpoint inhibitor toxicities. Trends in Immunology 46, 61–73 (2025). 10.1016/j.it.2024.11.014

57 Nolz, J. C. & Richer, M. J. Control of memory CD8(+) T cell longevity and effector functions by IL-15. Mol Immunol 117, 180–188 (2020). 10.1016/j.molimm.2019.11.011

58 Bruchfeld, A., Wendt, M. & Miller, E. J. Macrophage Migration Inhibitory Factor in Clinical Kidney Disease. Front Immunol 7, 8 (2016). 10.3389/fimmu.2016.00008

59 Ding, N., Li, P.-L., Wu, K.-L., Lv, T.-G., Yu, W.-L. & Hao, J. Macrophage migration inhibitory factor levels are associated with disease activity and possible complications in membranous nephropathy. Scientific Reports 12, 18558 (2022). 10.1038/s41598-022-23440-1

60 Kong, Y. Z., Chen, Q. & Lan, H. Y. Macrophage Migration Inhibitory Factor (MIF) as a Stress Molecule in Renal Inflammation. Int J Mol Sci 23 (2022). 10.3390/ijms23094908

61 Mu, Y.-F. et al. Macrophage-driven inflammation in acute kidney injury: Therapeutic opportunities and challenges. Translational Research 278, 1–9 (2025). 10.1016/j.trsl.2025.02.003

62 Murakami, N., Borges, T. J., Yamashita, M. & Riella, L. V. Severe acute interstitial nephritis after combination immune-checkpoint inhibitor therapy for metastatic melanoma. Clin Kidney J 9, 411–417 (2016). 10.1093/ckj/sfw024

63 Tibbs, E. & Cao, X. Emerging Canonical and Non-Canonical Roles of Granzyme B in Health and Disease. Cancers (Basel) 14 (2022). 10.3390/cancers14061436

64 Yarchoan, M. et al. Effects of B cell–activating factor on tumor immunity. JCI Insight 5 (2020). 10.1172/jci.insight.136417

65 Wu, J., Li, Z., Zheng, S., Liu, Y. & Zhao, N. BAFF in autoimmune kidney diseases: pathogenic pathways, biomarker potential, and therapeutic horizons. Ren Fail 47, 2577164 (2025). 10.1080/0886022x.2025.2577164

66 Inoue, Y. et al. Serum immune modulators associated with immune-related toxicities and efficacy of atezolizumab in patients with non-small cell lung cancer. J Cancer Res Clin Oncol 149, 2963–2974 (2023). 10.1007/s00432-022-04193-w

67 Mackay, F. & Browning, J. L. BAFF: A fundamental survival factor for B cells. Nature Reviews Immunology 2, 465–475 (2002). 10.1038/nri844

68 Dhodapkar, K. M., Duffy, A. & Dhodapkar, M. V. Role of B cells in immune-related adverse events following checkpoint blockade. Immunol Rev 318, 89–95 (2023). 10.1111/imr.13238

69 Yarchoan, M. et al. Effects of B cell-activating factor on tumor immunity. JCI Insight 5 (2020). 10.1172/jci.insight.136417

70 Mistry, K., Sadarangani, S., Moreno, D., Mejia, S. M., Moledina, D. G. & Sise, M. E. Novel Biomarkers and Imaging Tests for Acute Kidney Injury Diagnosis in Patients with Cancer. Kidne*y360* 6 (2025).

71 Tang, J., Liu, N. & Zhuang, S. Role of epidermal growth factor receptor in acute and chronic kidney injury. Kidney Int 83, 804–810 (2013). 10.1038/ki.2012.435

72 Zeng, F., Singh, A. B. & Harris, R. C. The role of the EGF family of ligands and receptors in renal development, physiology and pathophysiology. Exp Cell Res 315, 602–610 (2009). 10.1016/j.yexcr.2008.08.005

73 Gupta, S. et al. Urinary C-X-C-motif ligand 9 (CXCL9) in immune checkpoint inhibitor-associated acute interstitial nephritis. Kidney Int 108, 491–496 (2025). 10.1016/j.kint.2025.05.029

74 Baek, J.-H. et al. IL-34 mediates acute kidney injury and worsens subsequent chronic kidney disease. The Journal of Clinical Investigation 125, 3198–3214 (2015). 10.1172/JCI81166

75 Morel, D. S. et al. RENAL SYNTHESIS OF LEUKAEMIA INHIBITORY FACTOR (LIF), UNDER NORMAL AND INFLAMMATORY CONDITIONS. Cytokine 12, 265–271 (2000). 10.1006/cyto.1999.0545

76 van Ham, S. M. et al. Urinary granzyme A mRNA is a biomarker to diagnose subclinical and acute cellular rejection in kidney transplant recipients. Kidney International 78, 1033–1040 (2010). 10.1038/ki.2010.274

77 Kummer, J. A., Wever, P. C., Kamp, A. M., ten Berge, I. J., Hack, C. E. & Weening, J. J. Expression of granzyme A and B proteins by cytotoxic lymphocytes involved in acute renal allograft rejection. Kidney Int 47, 70–77 (1995). 10.1038/ki.1995.8

78 Kim, S. T. et al. Distinct molecular and immune hallmarks of inflammatory arthritis induced by immune checkpoint inhibitors for cancer therapy. Nature Communications 13, 1970 (2022). 10.1038/s41467-022-29539-3

79 Wang, S. J., Dougan, S. K. & Dougan, M. Immune mechanisms of toxicity from checkpoint inhibitors. Trends Cancer 9, 543–553 (2023). 10.1016/j.trecan.2023.04.002

80 Irazabal, M. V. & Torres, V. E. Reactive Oxygen Species and Redox Signaling in Chronic Kidney Disease. Cells 9 (2020). 10.3390/cells9061342

81 McWilliam, S. J. et al. The complex interplay between kidney injury and inflammation. Clinical Kidney Journal 14, 780–788 (2021). 10.1093/ckj/sfaa164

82 Lv, W., Booz, G. W., Wang, Y., Fan, F. & Roman, R. J. Inflammation and renal fibrosis: Recent developments on key signaling molecules as potential therapeutic targets. Eur J Pharmacol 820, 65–76 (2018). 10.1016/j.ejphar.2017.12.016

83 Meng, X.-m., Wang, L., Nikolic-Paterson, D. J. & Lan, H.-Y. Innate immune cells in acute and chronic kidney disease. Nature Reviews Nephrology 21, 464–482 (2025). 10.1038/s41581-025-00958-x

84 Liu, Q., Zhu, W., Tang, C., Liu, W. & Luo, X. Integrative analysis of ASXL family genes reveals ASXL2 as an immunoregulatory molecule in head and neck squamous cell carcinoma. Scientific Reports 14, 31368 (2024). 10.1038/s41598-024-82815-8

85 Jin, J. et al. Role of epigenetically regulated inflammation in renal diseases. Seminars in Cell & Developmental Biology 154, 295–304 (2024). 10.1016/j.semcdb.2022.10.005

86 Miao, J., Sise, M. E. & Herrmann, S. M. Immune checkpoint inhibitor related nephrotoxicity: Advances in clinicopathologic features, noninvasive approaches, and therapeutic strategy and rechallenge. Front Nephrol 2, 1017921 (2022). 10.3389/fneph.2022.1017921

87 Francisco, L. M., Sage, P. T. & Sharpe, A. H. The PD-1 pathway in tolerance and autoimmunity. Immunol Rev 236, 219–242 (2010). 10.1111/j.1600-065X.2010.00923.x

88 Gomez-Preciado, F. et al. Urinary soluble PD-1 as a biomarker of checkpoint inhibitor-induced acute tubulointerstitial nephritis. Clin Kidney J 17, sfae200 (2024). 10.1093/ckj/sfae200

89 Wei, Y. & Jiang, Z. The role of programed death-ligand 1 in renal diseases. Journal of Receptors and Signal Transduction 40, 295–300 (2020). 10.1080/10799893.2020.1734820

90 Pippin, J. W. et al. Upregulated PD-1 signaling antagonizes glomerular health in aged kidneys and disease. The Journal of Clinical Investigation 132 (2022).

91 Curran, C. S., Gupta, S., Sanz, I. & Sharon, E. PD-1 immunobiology in systemic lupus erythematosus. J Autoimmun 97, 1–9 (2019). 10.1016/j.jaut.2018.10.025

92 Zhang, S., Zhao, J., Bai, X., Handley, M. & Shan, F. Biological effects of IL-15 on immune cells and its potential for the treatment of cancer. International Immunopharmacology 91, 107318 (2021). 10.1016/j.intimp.2020.107318

93 Steel, J. C., Waldmann, T. A. & Morris, J. C. Interleukin-15 biology and its therapeutic implications in cancer. Trends Pharmacol Sci 33, 35–41 (2012). 10.1016/j.tips.2011.09.004

94 Correia, M. P., Costa, A. V., Uhrberg, M., Cardoso, E. M. & Arosa, F. A. IL-15 induces CD8+ T cells to acquire functional NK receptors capable of modulating cytotoxicity and cytokine secretion. Immunobiology 216, 604–612 (2011). 10.1016/j.imbio.2010.09.012

95 Franz, T., Negele, J. & Kahlfuss, S. Cytotoxic innate lymphoid cells sense tumor-derived IL-15: a novel mechanism of cancer immunosurveillance. Signal Transduction and Targeted Therapy 7, 326 (2022). 10.1038/s41392-022-01173-x

96 Skariah, N., James, O. J. & Swamy, M. Signalling mechanisms driving homeostatic and inflammatory effects of interleukin-15 on tissue lymphocytes. Discov Immunol 3, kyae002 (2024). 10.1093/discim/kyae002

97 Jabri, B. & Abadie, V. IL-15 functions as a danger signal to regulate tissue-resident T cells and tissue destruction. Nat Rev Immunol 15, 771–783 (2015). 10.1038/nri3919

98 Kang, L. et al. CCR8 Signaling via CCL1 Regulates Responses of Intestinal IFN-γ Producing Innate Lymphoid CelIs and Protects From Experimental Colitis. Front Immunol 11, 609400 (2020). 10.3389/fimmu.2020.609400

99 Percie du Sert, N., et al. The ARRIVE guidelines 2.0: Updated guidelines for reporting animal research. PLOS Biology 18, e3000410 (2020). 10.1371/journal.pbio.3000410

