## Supplemental for "Nephrotoxicity of Immune Checkpoint Inhibitors in Mice with a Human Immune System"

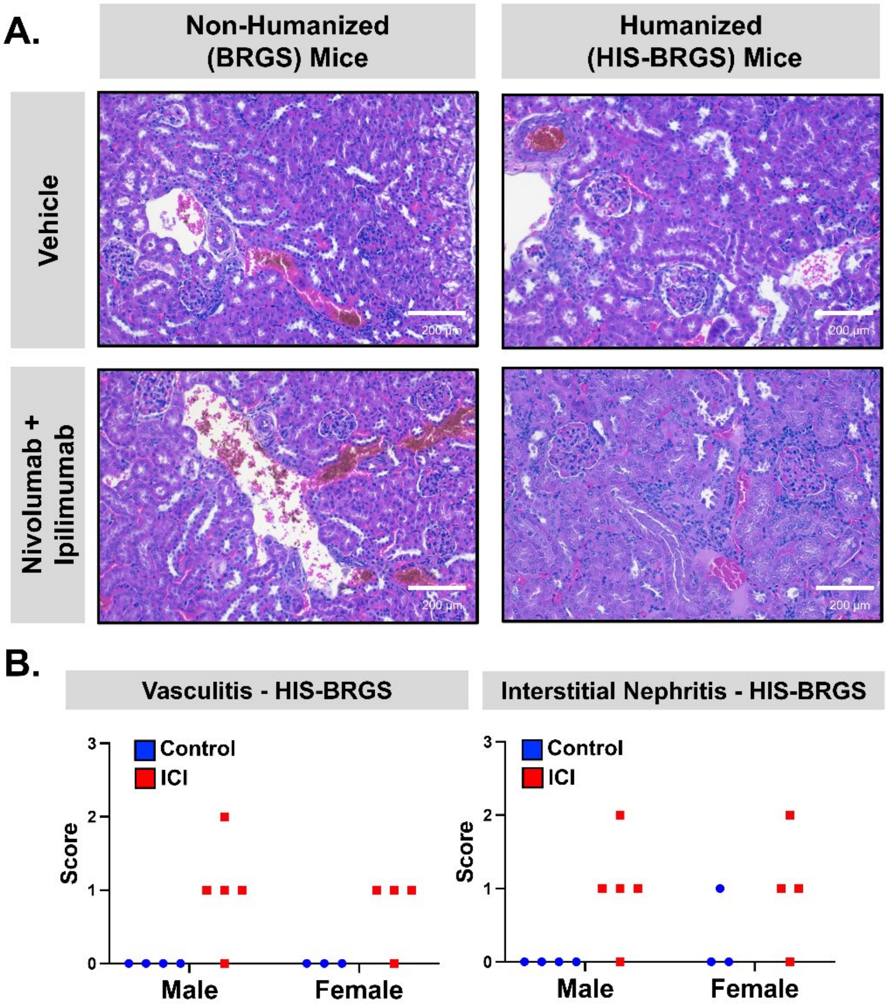


**Supplemental Figure 1. Kidney Histopathology By Animal Sex.** Humanized HIS-BRGS and non-humanized BRGS mice were implanted with tumors, followed by treatment with vehicle control (PBS) or immune checkpoint inhibitors (ICI, nivolumab and ipilimumab, 20 and 10 mg/kg weekly, i.p.). Kidneys were fixed in formalin and embedded in paraffin before sectioning and staining with H&E. (A) Light microscope images were acquired at 10X magnification. (B) Vasculitis and interstitial nephritis were assessed according to a range of scores (no lesions = 0; minimal lesions, <10% kidney affected = 1; mild lesions, 10-25% kidney affected = 2; moderate lesions, >25-40% kidney affected = 3) by a veterinary anatomic pathologist and displayed according to animal sex.

**Supplemental Table 1. Flow Cytometry Antibodies and Reagents**

| Flow Cytometry Antibodies | | | | | Flow Cytometry Reagents | |
|---|---|---|---|---|---|---|
| **Target** | **Species** | **Fluorochrome** | **Clone** | **Source** | **Reagent** | **Source** |
| **CD45** | anti-human | BUV395 | HI30 | BD Bioscience | Zombie Green | BioLegend |
| **CD5** | anti-human | Fitc | UCHT2 | BioLegend | L/D Blue | ThermoFisher |
| **CD3** | anti-human | PE | HIT3a | BioLegend | Rec Hu IL6 | R&D Systems |
| **CD11b** | anti-human | PE | ICRF44 | BioLegend | Rec Hu SCF | R&D Systems |
| **CD14** | anti-human | PE | 63D3 | BioLegend | Rec Hu FLT3L | R&D Systems |
| **CD33** | anti-human | PE | P67.6 | BioLegend | FCS | Gibco |
| **PD-1** | anti-human | PE | EH12.2H7 | BioLegend | FCS CD34+ media | Stemcell |
| **CD3** | anti-human | PeCy7 | HIT3a | BioLegend | HBSS | Gibco |
| **CD45** | anti-human | BUV395 | HI30 | BD Bioscience | IMDM | Gibco |
| **CD45** | anti-human | PeCy7 | HI30 | BioLegend | Golgi Stop | BD Bioscience |
| **IFNγ** | anti-human | PeCy7 | 4S.B3 | BioLegend | Cell Stim Cocktail | Invitrogen |
| **FoxP3** | anti-human | PacB | 259D | BioLegend | Saponin | Sigma |
| **CD45** | anti-human | BV421 | HI30 | BioLegend | BSA (Bovine serum Albumin) | Sigma |
| **CD11c** | anti-human | BV421 | s-hcl-3 | BioLegend | Formaldehyde | Fisher |
| **HLA-DR** | anti-human | BV480 | G48-6 | BD Bioscience | Transcription Factor Staining Buffer Set | eBioscience |
| **Granzyme B** | anti-human | BV510 | GB11 | BioLegend | | |
| **CD45RA** | anti-human | BV605 | HI100 | BioLegend | | |
| **CD197/ CCR7** | anti-human | BV650 | GO43H7 | BioLegend | | |
| **CD8** | anti-human | BV711 | RPA-T8 | BioLegend | | |
| **CD4** | anti-human | BV785 | RPA-T4 | BioLegend | | |
| **TIGIT** | anti-human | Kiravia Blue 520 | A15153G | BioLegend | | |
| **CD56** | anti-human | APC | HD56 | BioLegend | | |
| **CD19** | anti-human | APC | HIB19 | BioLegend | | |
| **TNFα** | anti-human | APC | MAb11 | BioLegend | | |
| **CD34** | anti-human | APC | | Biolegend | | |
| **CD25** | anti-human | Spark NIR685 | M-A251 | BioLegend | | |
| **PD-1** | anti-human | AF700 | EH12.2H7 | BioLegend | | |
| **CD8** | anti-human | APCFire | RPA-T8 | BioLegend | | |
| **mCD45** | anti-mouse | APCCY7 | 30-F11 | BioLegend | | |
| **CD19** | anti-human | APCFire810 | HIB19 | BioLegend | | |
| **FCR Block** | human | | | Miltenyi | | |
| **CD32** | mouse | BUV395 | 2.4G2 | BioLegend | | |

### Supplemental Table 2. Limits of Detection (LOD) for Analytes Used for Multiplex

| Human Immune Response Panel | | | | | |
|---|---|---|---|---|---|
| **Analyte** | **LOD** | **Analyte** | **LOD** | **Analyte** | **LOD** |
| **April** | 185 | **CXCL-6** | 0.7 | **IL-15** | 3.1 |
| **BAFF** | 2.3 | **CXCL-8** | 2.7 | **IL-16** | 15.7 |
| **CCL1** | 1.1 | **CXCL-9** | 10 | **IL-17/CTLA-8** | 3.8 |
| **CCL2** | 2.6 | **CXCL-10** | 2.2 | **IL-18** | 9.7 |
| **CCL3** | 3.1 | **CXCL-11** | 12.1 | **IL-20** | 13.9 |
| **CCL4** | 9.1 | **CXCL-13** | 15.2 | **IL-21** | 10.1 |
| **CCL7** | 4.4 | **FGF-2** | 10.2 | **IL-22** | 17.5 |
| **CCL8** | 1 | **Galectin-3** | 1007.6 | **IL-23** | 15.2 |
| **CCL11** | 2 | **GM-CSF** | 13.6 | **Il-27** | 10.3 |
| **CCL13** | 15.2 | **Granzyme-A** | 7.7 | **IL-31** | 16.1 |
| **CCL17** | 0.7 | **Granzyme-B** | 6.4 | **IL-34** | 10.7 |
| **CCL19** | 48.3 | **HGF** | 1.1 | **IL-37** | 2.2 |
| **CCL20** | 11.3 | **IFN alpha** | 8.9 | **LIF** | 4.6 |
| **CCL21** | 14 | **IFN gamma** | 10.7 | **M-CSF** | 16.1 |
| **CCL22/MDC** | 19.3 | **IL-1 alpha** | 1.9 | **MIF** | 0.9 |
| **CCL23** | 34.3 | **IL-1 beta** | 7 | **MMP-1** | 4.2 |
| **CCL24** | 5.4 | **IL-2** | 10 | **NGF-β** | 5.9 |
| **CCL25** | 10.2 | **IL-2R** | 100.4 | **PTX-3** | 29.4 |
| **CCL26** | 1.6 | **IL-3** | 31.8 | **SCF** | 4.5 |
| **CD30** | 8.5 | **IL-4** | 13 | **TNF alpha** | 14.6 |
| **CD40L** | 12.4 | **IL-5** | 10.2 | **TNF beta** | 0.7 |
| **CSF-3** | 14 | **IL-6** | 12.6 | **TNF-RII** | 2.9 |
| **CX3CL1** | 3.5 | **IL-7** | 1 | **TRAIL** | 3.4 |
| **CXCL1** | 2.8 | **IL-9** | 0.9 | **TREM-1** | 474 |
| **CXCL2** | 1 | **IL-10** | 4.2 | **TSLP** | 3.9 |
| **CXCL5** | 8 | **IL-12p70** | 7.2 | **TWEAK** | 137.7 |
| | | **IL-13** | 4.7 | **VEGF alpha** | 6 |
| **Mouse Kidney Injury Biomarker Panels** | | | | | |
| **Analyte** | **LOD** | **Analyte** | **LOD** | **Analyte** | **LOD** |
| **B2-Microglobulin** | 50 | **IP-10** | 5 | **Renin** | 50 |
| **Clusterin** | 200 | **KIM-1** | 10 | **TIMP-1** | 20 |
| **Cystatin C** | 50 | **NGAL** | 5 | **VEGF** | 1 |
| **EGF** | 40 | **OPN** | 10 | | |
| **Human Immune-Oncology Checkpoint Protein Panel** | | | | | |
| **Analyte** | **LOD** | **Analyte** | **LOD** | **Analyte** | **LOD** |
| **CD80** | 11.2 | **CTLA-4** | 9.3 | **PD-L1** | 1.3 |
| **CD86** | 86.1 | **PD-1** | 13.7 | | |

**Supplemental Table 3. Primers for qPCR Analysis of Human or mouse Genes.**

| Target | 5' → 3' | 3' → 5' |
|---|---|---|
| **Human** | | |
| **AK2** | TCCTACCACGAGGAGTTCAACC | TGGTAGGCTTGCAGGCGGATTT |
| **ASXL2** | GGACAGAATCCAGGTGCGAAAAG | GATGGAGACTGGAAAACGAGCC |
| **CCS** | CAGAATGGAGGATGAGCAGCTG | GAGCGTGCAATGATGCCACAGG |
| **CCR5** | TCTCTTCTGGGCTCCCTACAAC | CCAAGAGTCTCTGTCACCTGCA |
| **CD226** | GGTGATACAGGTGGTTCAGTCAG | GGCTGGATCTTTTCCCACCTCA |
| **CD3D** | GCATCACATGGGTAGAGGGA | ACAGCTCTGGCACATTCGAT |
| **CD3E** | CGGTGGCCACAATTGTCATA | TTTCCGGATGGCTCATAGT |
| **CD3G** | TCCTTGCTGTTGGGGTCTAC | GGAGAACACCTGGACTACTCTG |
| **CD5** | CGAGTTCTTGCCCTCCTTTGCT | TCCTGGCTGAAGAGCTGTCACA |
| **CD45** | TTCCTCGTCTGATAAGACAACAGT | TCTTTGCTGTAGTCAATCCAGTG |
| **CD84** | CTTCCAGACTCCTGAGGACCAA | ACATAGCCAGCACGCTCAGCAA |
| **CD8A** | TGCAACCACAGGAACCGA | TGCTCCCTCAAAAGGAAGGAT |
| **CD8B** | CCGGAAGACAGTGGCATCTA | TACAAAGTGGGCCCTTCTGG |
| **CXCL10** | AGTGGCATTCAAGGAGTACCT | TGATGGCCTTCGATTCTGGA |
| **CXCL13** | TATCCCTAGACGCTTCATTGATCG | CCATTCAGCTTGAGGGTCCACA |
| **GZMA** | CCACACGCGAAGGTGACCTTAA | CCTGCAACTTGGCACATGGTTC |
| **GZMB** | CGACAGTACCATTGAGTTGTGCG | TTCGTCCATAGGAGACAATGCCC |
| **GZMK** | TCCAGTATGGCGGACATCACGT | CGCCTAAAACCACAGTGGGAGA |
| **IL-15** | TTTGGGCTGTTTCAGTGCAG | ACTTTGCAACTGGGGTGAAC |
| **IL-16** | CAAGTCTCTCAAGGGGACCA | TGTGGCCTCTGCTGTAGATT |
| **ICOS** | CCCATAGGATGTGCAGCCTTTG | GGCTGTGTTCACTGCTCTCATG |
| **IL7R** | ATCGCAGCACTCACTGACCTGT | TCAGGCACTTTACCTCCACGAG |
| **LNPEP** | CAATGGCGGATAGAAAGCTGGTG | GCCATAGGTCATTCCACCACTTC |
| **OPTN** | ACTCTGACCAGCAGGCTTACCT | CTATGTCAGGCAGAACCTCTCC |
| **PGM2** | CGGATGCTGATAGACTTGCTGTG | TGAGAGCACTGCGATCCTGGTT |
| **PIK3R1** | CGCCTCTTCTTATCAAGCTCGTG | GAAGCTGTCGTAATTCTGCCAGG |
| **SAT2** | CTGAGAGCAGATGGCTTTGGAG | CCTCCAGATAAATGGTGCGTCC |
| **SHC1** | ACAGCCGAGTATGTCGCCTATG | CAATGGTGCTGATGACATCCTGG |
| **SLC12A4** | CTGTAACAGGCATCATGGCTGG | CACACTGCTGAAGTACACGAGG |
| **TAGAP** | ATGACTCCCTGGAGCACACTGA | CTGTTGGATTCCACATCAGGGTC |
| **Mouse** | | |
| **Gapdh** | GTTCCTACCCCCAATGTGTC | GTTGAAGTCGCAGGAGACAA |

**Supplemental Table 4. mRNA Expression of Podocyte Essential Genes in Vehicle-Treated BRGS and HIS-BRGS Mice.**[1]

| | Veh BRGS | | Veh HIS-BRGS | | |
|---|---|---|---|---|---|
| Target | Mean | SE | Mean | SE | P-adj |
| Aox1 | 0.57 | 0.38 | 0.16 | 0.03 | 0.0792 |
| Cd59a | 9.24 | 0.41 | 6.32 | 0.35 | 0.0044 |
| Epb41l5 | 8.50 | 0.32 | 6.34 | 0.20 | 0.0006 |
| Ezr | 131.49 | 7.27 | 101.20 | 2.74 | 0.0018 |
| Fnbp1l | 7.43 | 0.31 | 6.28 | 0.15 | 0.0532 |
| Golim4 | 3.64 | 0.07 | 2.76 | 0.10 | 0.0052 |
| Ift80 | 1.46 | 0.08 | 1.09 | 0.05 | 0.0195 |
| Itgav | 18.63 | 1.92 | 14.36 | 0.80 | 0.0774 |
| Itgb5 | 32.59 | 1.41 | 40.12 | 1.55 | 0.0284 |
| Lpl | 64.01 | 5.65 | 24.55 | 1.53 | 0.0000 |
| Magi2 | 1.41 | 0.08 | 1.09 | 0.06 | 0.0583 |
| Mtss1 | 12.33 | 0.54 | 6.87 | 0.43 | 0.0000 |
| Myom2 | 2.33 | 0.21 | 1.10 | 0.10 | 0.0000 |
| Nebl | 0.64 | 0.02 | 0.40 | 0.04 | 0.0166 |
| Nupr1 | 3.47 | 0.46 | 5.29 | 0.82 | 0.1703 |
| Plce1 | 1.33 | 0.02 | 0.91 | 0.07 | 0.0156 |
| Podxl | 54.62 | 3.86 | 36.67 | 2.86 | 0.0091 |
| Robo2 | 1.21 | 0.12 | 0.73 | 0.09 | 0.0441 |
| Sema3g | 24.98 | 1.22 | 20.02 | 1.28 | 0.1682 |
| Shisa3 | 4.26 | 0.52 | 2.10 | 0.18 | 0.0000 |
| Synpo | 8.85 | 0.15 | 5.55 | 0.34 | 0.0002 |
| Tdrd5 | 0.89 | 0.07 | 0.65 | 0.04 | 0.0963 |
| Thsd7a | 2.09 | 0.13 | 1.05 | 0.08 | 0.0000 |
| Wt1 | 2.01 | 0.09 | 1.13 | 0.09 | 0.0002 |

[1] Bulk RNA Sequencing was performed on total RNA from kidneys of vehicle-treated BRGS (n=3) and HIS-BRGS mice (n=7). Podocyte-essential genes were selected from Lu et al. and expressed as fragments per kilobase of transcript per million (FPKM) values and reported as mean and standard error (SE).

**Supplemental Table 5. Circulating Human Cytokines, Growth Factors, Chemokines, Proteases, and Soluble Receptors in Vehicle- and ICI-Treated BRGS and HIS-BRGS Mice.**[1]

| | Veh BRGS | | ICI BRGS | | Veh HIS-BRGS | | ICI HIS-BRGS | |
|---|---|---|---|---|---|---|---|---|
| **Target** | **Mean** | **SE** | **Mean** | **SE** | **Mean** | **SE** | **Mean** | **SE** |
| **BAFF** | 1.65 | 0.00 | 1.65 | 0.00 | 43.10 | 13.24 | 94.69 | 19.42 |
| **CCL1** | 0.77 | 0.00 | 0.77 | 0.00 | 73.66 | 29.59 | 162.66 | 39.17 |
| **CCL17** | 0.48 | 0.00 | 0.48 | 0.00 | 10.03 | 5.24 | 24.94 | 6.40 |
| **CCL20** | 18.05 | 6.93 | 16.91 | 8.92 | 18.02 | 8.68 | 10.33 | 1.83 |
| **CCL22/MDC** | 13.65 | 0.00 | 13.65 | 0.00 | 67.93 | 34.10 | 97.97 | 26.54 |
| **CCL24** | 3.82 | 0.00 | 3.82 | 0.00 | 46.33 | 40.11 | 35.26 | 13.91 |
| **CCL25** | 4.41 | 1.64 | 5.86 | 1.35 | 7.21 | 1.59 | 20.60 | 4.35 |
| **CCL7** | 3.10 | 0.00 | 5.54 | 2.44 | 7.15 | 2.54 | 14.33 | 1.34 |
| **CD30** | 6.04 | 0.00 | 6.04 | 0.00 | 422.53 | 143.33 | 2666.99 | 880.30 |
| **CSF-3** | 10.82 | 0.92 | 9.90 | 0.00 | 14.59 | 4.11 | 18.52 | 4.13 |
| **CXCL1** | 3.17 | 0.82 | 2.32 | 0.21 | 14.58 | 4.12 | 14.61 | 2.93 |
| **CXCL10** | 1.58 | 0.00 | 1.58 | 0.00 | 97.60 | 74.01 | 146.82 | 53.07 |
| **CXCL11** | 8.56 | 0.00 | 8.56 | 0.00 | 19.47 | 7.07 | 32.18 | 12.95 |
| **CXCL8** | 15.02 | 12.17 | 12.82 | 3.72 | 78.25 | 36.48 | 125.61 | 43.39 |
| **CXCL13** | 10.72 | 0.00 | 10.72 | 0.00 | 2113.44 | 801.92 | 2625.73 | 515.09 |
| **FGF-2** | 146.23 | 91.32 | 69.44 | 21.21 | 195.53 | 119.74 | 223.70 | 25.58 |
| **Galectin-3** | 712.48 | 0.00 | 712.48 | 0.00 | 747.94 | 35.46 | 17306.94 | 9441.35 |
| **GM-CSF** | 9.62 | 0.00 | 9.62 | 0.00 | 41.78 | 32.16 | 145.74 | 76.28 |
| **Granzyme-A** | 5.44 | 0.00 | 5.44 | 0.00 | 7.29 | 3.20 | 33.20 | 20.94 |
| **HGF** | 0.60 | 0.17 | 0.72 | 0.18 | 0.83 | 0.30 | 1.70 | 0.40 |
| **IL-2R** | 70.99 | 0.00 | 70.99 | 0.00 | 895.26 | 697.36 | 4849.32 | 2415.56 |
| **IL-3** | 22.49 | 0.00 | 22.49 | 0.00 | 23.45 | 0.97 | 22.49 | 0.00 |
| **IL-6** | 8.89 | 0.00 | 8.89 | 0.00 | 154.80 | 115.64 | 213.72 | 118.85 |
| **IL-15** | 2.21 | 0.00 | 2.21 | 0.00 | 28.43 | 9.20 | 56.36 | 12.75 |
| **IL-16** | 11.10 | 0.00 | 11.10 | 0.00 | 61.26 | 31.03 | 381.74 | 149.74 |
| **IL-18** | 14.43 | 8.39 | 7.81 | 0.95 | 46.82 | 22.89 | 52.14 | 15.09 |
| **IL-34** | 10.37 | 2.83 | 7.54 | 0.00 | 8.98 | 2.53 | 21.23 | 4.44 |
| **LIF** | 224.43 | 30.92 | 203.46 | 23.65 | 118.87 | 20.24 | 218.16 | 13.82 |
| **MIF** | 37.14 | 7.91 | 24.59 | 3.41 | 18.75 | 4.74 | 36.21 | 6.29 |
| **MMP-1** | 8.05 | 5.64 | 12.22 | 3.87 | 78.38 | 30.53 | 102.95 | 60.49 |
| **PTX-3** | 349.27 | 316.47 | 65.19 | 25.47 | 54.50 | 9.90 | 88.67 | 37.78 |
| **TNF-RII** | 2.07 | 0.00 | 2.07 | 0.00 | 25.79 | 12.88 | 67.69 | 23.56 |

[1]Proteins below the LOD: APRIL, CCL-2, CCL-3, CCL-4, CCL-8, CCL-11, CCL-26, CCL-13, CCL-19, CCL-21, CCL-23, CD40L, CTLA-8, CXCL-2, CXCL-5, CXCL-6, CXCL-9, CX3CL1, Granzyme B, IFN alpha, IFN gamma, IL-1 alpha, IL-1 beta, IL-2, IL-3, IL-4, IL-5, IL-7, IL-9, IL-10, IL-12p70, IL-13, IL-20, IL-21, IL-22, IL-23, IL-27, IL-31, IL-37, M-CSF, NGF-β, SCF, TNF alpha, TNF beta, TRAIL, TREM1, TSLP, TWEAK, VEGF alpha

**Supplemental Table 6. Human Cytokines, Growth Factors, Chemokines, Proteases, and Soluble Receptors in the Kidneys of Vehicle- and ICI-Treated BRGS and HIS-BRGS Mice.**

| | Veh BRGS | | ICI BRGS | | Veh HIS-BRGS | | ICI HIS-BRGS | |
|---|---|---|---|---|---|---|---|---|
| **Target** | **Mean** | **SE** | **Mean** | **SE** | **Mean** | **SE** | **Mean** | **SE** |
| **April** | 561.01 | 20.00 | 559.07 | 19.52 | 632.11 | 84.24 | 627.87 | 103.35 |
| **BAFF** | 3.25 | 0.08 | 3.26 | 0.07 | 6.23 | 0.55 | 11.22 | 0.95 |
| **CCL1** | 0.85 | 0.02 | 0.81 | 0.03 | 8.87 | 2.17 | 32.40 | 6.34 |
| **CCL2** | 0.74 | 0.09 | 0.67 | 0.06 | 4.65 | 2.09 | 2.51 | 0.75 |
| **CCL3** | 1.15 | 0.16 | 1.18 | 0.10 | 1.82 | 0.28 | 4.33 | 2.07 |
| **CCL4** | 4.96 | 0.61 | 4.93 | 0.35 | 6.27 | 0.56 | 12.24 | 3.94 |
| **CCL7** | 11.48 | 0.18 | 11.49 | 0.21 | 12.91 | 0.65 | 13.61 | 0.84 |
| **CCL8** | 0.34 | 0.04 | 0.35 | 0.02 | 0.43 | 0.04 | 0.37 | 0.03 |
| **CCL11** | 0.99 | 0.08 | 1.01 | 0.04 | 1.02 | 0.06 | 0.98 | 0.10 |
| **CCL13** | 16.22 | 1.63 | 16.22 | 0.84 | 27.33 | 5.94 | 18.90 | 2.16 |
| **CCL17** | 1.14 | 0.11 | 1.13 | 0.07 | 2.36 | 0.44 | 5.28 | 2.19 |
| **CCL19** | 124.50 | 8.65 | 129.30 | 6.33 | 152.87 | 15.32 | 167.99 | 36.03 |
| **CCL20** | 35.75 | 0.99 | 35.74 | 1.61 | 40.83 | 3.71 | 39.58 | 6.84 |
| **CCL21** | 31.61 | 1.44 | 33.75 | 1.42 | 47.85 | 7.62 | 49.76 | 12.22 |
| **CCL22/MDC** | 10.45 | 1.19 | 10.47 | 0.58 | 13.34 | 1.35 | 23.45 | 7.46 |
| **CCL23** | 22.47 | 0.94 | 22.24 | 0.66 | 38.56 | 4.43 | 36.99 | 5.92 |
| **CCL24** | 3.35 | 0.31 | 3.39 | 0.21 | 4.10 | 0.42 | 3.99 | 0.34 |
| **CCL25** | 8.54 | 0.92 | 9.09 | 0.68 | 10.69 | 1.05 | 11.07 | 1.42 |
| **CCL26** | 1.66 | 0.13 | 1.67 | 0.10 | 2.62 | 0.32 | 2.79 | 0.46 |
| **CD30** | 4.29 | 0.39 | 4.52 | 0.23 | 5.90 | 0.55 | 11.80 | 2.71 |
| **CD40L** | 10.62 | 0.77 | 10.99 | 0.70 | 17.71 | 2.38 | 17.81 | 3.55 |
| **CSF-3** | 70.82 | 11.73 | 70.30 | 6.70 | 77.38 | 10.63 | 69.22 | 8.18 |
| **CX3CL1** | 5.25 | 0.17 | 5.23 | 0.20 | 5.85 | 0.62 | 5.80 | 0.75 |
| **CXCL1** | 1.16 | 0.12 | 1.18 | 0.10 | 1.92 | 0.22 | 1.71 | 0.10 |
| **CXCL2** | 2.03 | 0.10 | 2.23 | 0.09 | 2.79 | 0.30 | 2.87 | 0.44 |
| **CXCL5** | 1.40 | 0.36 | 1.22 | 0.22 | 4.12 | 1.57 | 2.52 | 0.37 |
| **CXCL6** | 1.03 | 0.08 | 1.07 | 0.05 | 1.22 | 0.10 | 1.19 | 0.17 |
| **CXCL8** | 16.22 | 4.46 | 14.91 | 3.64 | 28.77 | 4.50 | 36.48 | 7.14 |
| **CXCL-9** | 58.19 | 2.49 | 61.92 | 2.10 | 69.14 | 7.11 | 71.38 | 11.34 |
| **CXCL10** | 1.15 | 0.11 | 1.18 | 0.13 | 7.63 | 4.35 | 19.52 | 8.00 |
| **CXCL11** | 30.69 | 0.31 | 30.72 | 1.14 | 34.71 | 4.05 | 35.40 | 5.30 |
| **CXCL13** | 34.04 | 1.81 | 33.31 | 1.22 | 115.98 | 24.66 | 415.58 | 222.81 |
| **FGF-2** | 80.62 | 11.82 | 70.77 | 6.43 | 70.60 | 4.26 | 68.91 | 9.82 |
| **Galectin-3** | 1663.95 | 76.13 | 1558.82 | 40.51 | 1759.88 | 151.28 | 1742.75 | 200.75 |
| **GM-CSF** | 15.34 | 2.19 | 14.83 | 1.17 | 18.31 | 1.33 | 18.58 | 2.08 |
| **Granzyme-A** | 2.73 | 0.23 | 2.95 | 0.17 | 9.33 | 2.00 | 49.68 | 16.62 |
| **Granzyme-B** | 5.86 | 0.40 | 5.84 | 0.17 | 7.23 | 0.87 | 18.26 | 7.46 |
| **HGF** | 0.69 | 0.02 | 0.67 | 0.03 | 1.03 | 0.16 | 1.05 | 0.16 |
| **IFN alpha** | 0.63 | 0.00 | 0.63 | 0.00 | 1.22 | 0.43 | 1.34 | 0.46 |
| **IFN gamma** | 1.06 | 0.32 | 1.21 | 0.20 | 1.42 | 0.20 | 2.12 | 0.49 |
| **IL-1 alpha** | 2.49 | 0.36 | 2.57 | 0.22 | 3.22 | 0.35 | 3.22 | 0.44 |
| **IL-1 beta** | 1.69 | 0.28 | 1.70 | 0.13 | 2.52 | 0.34 | 2.50 | 0.36 |

| | | | | | | | | |
|---|---|---|---|---|---|---|---|---|
| **IL-2** | 3.25 | 0.42 | 3.37 | 0.23 | 3.52 | 0.24 | 3.12 | 0.39 |
| **IL-2R** | 63.75 | 4.74 | 54.80 | 10.08 | 123.09 | 19.33 | 181.68 | 55.80 |
| **IL-3** | 106.11 | 7.31 | 105.62 | 5.42 | 131.41 | 21.19 | 128.78 | 23.64 |
| **IL-4** | 7.08 | 2.02 | 10.99 | 1.16 | 22.39 | 3.91 | 11.61 | 2.63 |
| **IL-5** | 10.34 | 0.20 | 9.74 | 0.28 | 10.55 | 1.12 | 10.70 | 1.74 |
| **IL-6** | 8.66 | 1.40 | 7.82 | 0.79 | 23.34 | 8.47 | 30.28 | 12.03 |
| **IL-7** | 1.29 | 0.15 | 1.47 | 0.09 | 1.69 | 0.17 | 1.71 | 0.26 |
| **IL-9** | 0.76 | 0.06 | 0.81 | 0.05 | 0.83 | 0.07 | 0.83 | 0.10 |
| **IL-10** | 1.43 | 0.16 | 1.53 | 0.13 | 2.27 | 0.30 | 2.00 | 0.29 |
| **IL-13** | 1.16 | 0.03 | 0.94 | 0.11 | 1.13 | 0.17 | 1.14 | 0.14 |
| **IL-15** | 2.47 | 0.12 | 2.24 | 0.13 | 5.13 | 0.53 | 9.28 | 0.82 |
| **IL-16** | 7.56 | 1.14 | 7.80 | 0.67 | 12.36 | 2.22 | 123.08 | 63.37 |
| **IL-17/CTLA-8** | 2.20 | 0.24 | 2.24 | 0.14 | 2.61 | 0.24 | 2.73 | 0.26 |
| **IL-18** | 52.36 | 7.75 | 48.06 | 3.39 | 55.13 | 9.89 | 50.79 | 9.14 |
| **IL-20** | 9.14 | 1.14 | 9.71 | 0.60 | 10.05 | 0.80 | 9.65 | 0.90 |
| **IL-21** | 56.62 | 4.24 | 57.79 | 2.53 | 71.20 | 10.10 | 73.49 | 14.10 |
| **IL-22** | 43.43 | 1.32 | 43.99 | 2.01 | 53.70 | 8.73 | 52.65 | 10.72 |
| **IL-23** | 6.75 | 0.59 | 7.06 | 0.43 | 7.78 | 0.95 | 6.81 | 0.53 |
| **IL-27** | 1.77 | 0.62 | 1.99 | 0.46 | 5.71 | 0.19 | 4.51 | 1.65 |
| **IL-31** | 13.43 | 1.69 | 13.67 | 1.15 | 30.79 | 4.31 | 26.70 | 4.71 |
| **IL-34** | 9.72 | 1.24 | 9.08 | 0.64 | 13.47 | 2.51 | 13.79 | 2.73 |
| **IL-37** | 3.25 | 0.16 | 3.39 | 0.23 | 4.73 | 0.66 | 4.81 | 0.87 |
| **LIF** | 5.80 | 0.57 | 5.68 | 0.38 | 6.06 | 0.34 | 7.09 | 0.76 |
| **M-CSF** | 84.91 | 13.88 | 91.51 | 9.16 | 82.02 | 11.16 | 78.63 | 7.54 |
| **MIF** | 9.67 | 3.62 | 4.61 | 0.91 | 12.20 | 1.83 | 27.46 | 5.83 |
| **MMP-1** | 1.81 | 0.50 | 1.89 | 0.31 | 2.60 | 0.29 | 2.14 | 0.17 |
| **NGF-β** | 3.34 | 0.47 | 3.57 | 0.22 | 4.16 | 0.36 | 7.28 | 1.73 |
| **PTX-3** | 12.26 | 3.35 | 10.55 | 1.46 | 21.57 | 3.82 | 18.27 | 3.81 |
| **SCF** | 2.73 | 0.32 | 2.66 | 0.19 | 2.66 | 0.24 | 2.64 | 0.29 |
| **TNF alpha** | 0.67 | 0.13 | 0.65 | 0.12 | 1.65 | 0.34 | 1.84 | 0.38 |
| **TNF beta** | 0.49 | 0.02 | 0.45 | 0.03 | 0.61 | 0.09 | 0.61 | 0.12 |
| **TNF-RII** | 3.96 | 0.36 | 3.84 | 0.26 | 5.91 | 0.73 | 8.69 | 1.69 |
| **TRAIL** | 49.58 | 1.05 | 48.72 | 1.87 | 52.81 | 7.26 | 54.79 | 9.72 |
| **TREM-1** | 186.88 | 39.73 | 251.49 | 41.13 | 467.50 | 99.90 | 445.79 | 154.49 |
| **TSLP** | 8.50 | 0.91 | 8.89 | 0.63 | 14.06 | 1.80 | 14.04 | 2.45 |
| **TWEAK** | 66.92 | 10.57 | 55.76 | 4.89 | 80.81 | 4.71 | 92.67 | 8.00 |
| **VEGF alpha** | 19.55 | 4.39 | 24.32 | 4.08 | 25.64 | 1.77 | 22.17 | 3.50 |

[1]Proteins below the LOD: IL-12p70.
