## Supplementary material for "Nephrotoxicity of Immune Checkpoint Inhibitors in Mice with a Human Immune System": ARRIVE

### The ARRIVE reporting checklist

For checking that articles describing *in vivo* animal experiments can be understood and used by everyone

| | Item Description | Location (or reason for not reporting) |
|---|---|---|
| **Essential 10** | | |
| 1. Study Design | | |
| 1a. The groups being compared | For each experiment, describe the groups being compared, including control groups. If you did not use a control group, explain why. | Methods, Tumor-Bearing Human Immune System Mice; Results, Animal Characteristics |
| 1b. The experimental unit | Describe the experimental unit (e.g., a single animal, litter, or cage of animals). | Methods, Tumor-Bearing Human Immune System Mice; Results, Animal Characteristics |
| 2. Sample Size | | |
| 2a. Number of Experimental Units | Specify the exact number of experimental units allocated to each group, and the total number in each experiment. Also indicate the total number of animals used. | Methods, Tumor-Bearing Human Immune System Mice; Results, Animal Characteristics |
| 2b. Sample Size Justification | Explain how the sample size was decided. Provide details of any a priori sample size calculation, if done. | Methods, Tumor-Bearing Human Immune System Mice; Results, Animal Characteristics |
| 3. Inclusion and Exclusion Criteria | | |
| 3a. Inclusion and exclusion criteria | Describe any criteria used for including or excluding animals (or experimental units) during the experiment, and data points during the analysis. Specify if these criteria were established a priori. If no criteria were set, state this explicitly. | Methods, Tumor-Bearing Human Immune System Mice; Results, Animal Characteristics |
| 3b. Exclusions and Attritions | For each experimental group, report any animals, experimental units, or data points not included in the analysis and explain why. If there were no exclusions, state so. | Methods, Tumor-Bearing Human Immune System Mice; Results, Animal Characteristics |
| 3c. Numbers analysed | For each analysis, report the exact value of *n* in each experimental group. | Methods, Tumor-Bearing Human Immune System Mice; Results, Animal Characteristics |
| 4. Randomisation | | |
| 4a. Randomisation Use | State whether randomisation was used to allocate experimental units to control and treatment groups. If done, provide the method used to generate the randomisation sequence. | Methods, Tumor-Bearing Human Immune System Mice |

| | | |
|---|---|---|
| 4b. Confounders | Describe the strategy used to minimise potential confounders such as the order of treatments and measurements, or animal/cage location. If confounders were not controlled, state this explicitly. | Methods, Tumor-Bearing Human Immune System Mice, Results, Animal Characteristics Supplemental Figure 1 |
| 5. Blinding/Masking | Describe who was aware of the group allocation at the different stages of the experiment (during the allocation, the conduct of the experiment, the outcome assessment, and the data analysis). | Methods, Histopathology; no blinding required for experiment/analysis |
| 6. Outcome Measures | | |
| 6a. Outcome Measures | Clearly define all outcome measures assessed (e.g., cell death, molecular markers, or behavioural changes). | Methods - Histopathology; Transmission Electron Microscopy; Flow Cytometry; Multiplex ELISAs; qPCR and RNAseq |
| 6b. Primary Outcome Measure | For hypothesis-testing studies, specify the primary outcome measure, i.e., the outcome measure that was used to determine the sample size. | N/A - Methods, Histopathology |
| 7. Statistical Methods | | |
| 7a. Statistical Methods used for each Analysis | Provide details of the statistical methods used for each analysis, including software used. | Methods, Statistical Analysis |
| 7b. Statistical Assumptions | Describe any methods used to assess whether the data met the assumptions of the statistical approach, and what was done if the assumptions were not met. | Methods, Statistical Analysis |
| 8. Experimental Animals | | |
| 8a. Species-appropriate Details | Provide species-appropriate details of the animals used, including species, strain and substrain, sex, age or developmental stage, and, if relevant, weight. | Methods, Tumor-Bearing Human Immune System Mice; Results, Animal Characteristics Table 1; Supplemental Figure 1 |
| 8b. Further Information | Provide further relevant information on the provenance of animals, health/immune status, genetic modification status, genotype, and any previous procedures. | Methods, Tumor-Bearing Human Immune System Mice |
| 9. Experimental Procedures | | |
| 9a. What was done | What was done, how it was done, and what was used. | Methods, Tumor-Bearing Human Immune System Mice; Histopathology; Transmission Electron Microscopy; Flow |

| | | |
|---|---|---|
| | | Cytometry; Multiplex ELISAs; qPCR and RNAseq |
| 9b. When and how often procedures were conducted | For each experimental group, including controls, describe **when and how often** procedures were performed. | Methods, Tumor-Bearing Human Immune System Mice |
| 9c. Where procedures were conducted | For each experimental group, including controls, describe **where** procedures were conducted (including detail of any acclimatisation periods). | Acknowledgments; Methods, Tumor-Bearing Human Immune System Mice |
| 9d. Why procedures were done | For each experimental group, including controls, describe **why** procedures were conducted. | Introduction, para 3; Methods, Tumor-Bearing Human Immune System Mice |
| 10. Results | | |
| 10a. Summary/Descriptive Statistics per group | For each experiment conducted, including independent replications, report a summary/descriptive statistics for each experimental group, with a measure of variability where applicable (e.g., mean and SD, or median and range). | Methods, Statistical Analysis; Results throughout (Figures 1–6, Table 1) |
| 10b. Effect sizes and confidence intervals | For each experiment conducted, including independent replications, report the effect size with a confidence interval, if applicable. | Results throughout (Figures 1–6) |
| **Recommended Set** | | |
| 11. Abstract | Provide an accurate summary of the research objectives, animal species, strain and sex, key methods, principal findings, and study conclusions. | Abstract |
| 12. Background | | |
| 12a. Rationale | Include sufficient scientific background to understand the rationale and context for the study, and explain the experimental approach. | Introduction, paras 1–3 |
| 12b. Species and model | Explain how the animal species and model used address the scientific objectives and, where appropriate, the relevance to human biology. | Introduction, paras 2–3; Discussion, Elevated Baseline Immune Infiltration |
| 13. Objectives | Clearly describe the research question, research objectives and, where appropriate, specific hypotheses being tested. | Introduction, para 3; Abstract |
| 14. Ethical statement | Provide the name of the ethical review committee or equivalent that has approved the use of animals in this study and any relevant licence or protocol numbers (if applicable). If ethical approval was not sought or granted, provide a justification. | Methods, Tumor-Bearing Human Immune System Mice; Acknowledgments |
| 15. Housing and husbandry | Provide details of housing and husbandry conditions, including any environmental enrichment. | Acknowledgments |
| 16. Animal Care and Monitoring | | |

| | | |
|---|---|---|
| 16a. Reducing pain, suffering, and distress | Describe any interventions or steps taken in the experimental protocols to reduce pain, suffering, and distress. | Methods, Tumor-Bearing Human Immune System Mice; Acknowledgments |
| 16b. Adverse events | Report any expected or unexpected adverse events. | Results, Animal Characteristics |
| 16c. Humane endpoints | Describe the humane endpoints established for the study, the signs that were monitored, and the frequency of monitoring. If the study did not set humane endpoints, state this. | Methods, Tumor-Bearing Human Immune System Mice; Humane endpoints designated, approved, and monitored by CU Institutional Animal Care and Use Committe |
| 17. Interpretation/ scientific implications | | |
| 17a. Interpretation/scientific implications | Interpret the results, taking into account the study objectives and hypotheses, current theory, and other relevant studies in the literature. | Discussion |
| 17b. Limitations | Comment on the study limitations, including potential sources of bias, limitations of the animal model, and imprecision associated with the results. | Discussion, Elevated Baseline Immune Infiltration; Discussion, Checkpoint Molecule Induction |
| 18. Generalisability/translation | Comment on whether, and how, the findings of this study are likely to generalise to other species or experimental conditions, including any relevance to human biology (where appropriate). | Translational Statement; Discussion, Conclusions |
| 19. Protocol registration | Provide a statement indicating whether a protocol (including the research question, key design features, and analysis plan) was prepared before the study, and if and where this protocol was registered. | Methods, Tumor-Bearing Human Immune System Mice; |
| 20. Data Access | Provide a statement describing if and where study data are available. | Data Sharing Statement |
| 21. Declaration of interests | | |
| 21a. Conflicts of interests | Declare any potential conflicts of interest, including financial and nonfinancial. If none exist, this should be stated. | Disclosure |
| 21b. Funding | List all funding sources (including grant identifier) and the role of the funder(s) in the design, analysis, and reporting of the study. | Acknowledgments |
